# Phase separation as a tunable regulator of canonical gene regulatory motifs

**DOI:** 10.1101/2025.10.26.684618

**Authors:** Priya Chakraborty, Mintu Nandi, Sandeep Choubey

**Affiliations:** The Institute of Mathematical Sciences, CIT Campus, Taramani, Chennai 600113, India; Universal Biology Institute, The University of Tokyo, 7-3-1 Hongo, Bunkyo-ku, Tokyo 113-0033, Japan; Homi Bhabha National Institute, Training School Complex, Anushaktinagar, Mumbai 400094, India

## Abstract

Gene regulatory networks govern essential cellular processes such as signal transduction, metabolism, and cell fate control. Within these networks, canonical motifs such as genetic toggle switches and repressilators serve as building blocks that generate bistable and oscillatory behaviors. An open question is how cells regulate the dynamical behavior of such motifs, both in natural contexts and in synthetic systems. Recent studies highlight phase separation of transcription factors (TFs) and nucleic acids as a key organizing principle of intracellular biochemistry. In this work, we explore how phase separation of TFs influences the dynamics of toggle switch and repressilator using mean-field theory and stochastic simulations. Our mean-field analysis reveals that phase separation reshapes motif dynamics by altering fixed-point stability and basin geometry in toggle switch, and by altering oscillatory cycles in repressilator. A key finding for both motifs is that when multiple TFs undergo phase separation, the one with the lowest saturation concentration for phase separation dominates system dynamics. Interestingly, stochastic simulations show that the impact of phase separation on fluctuations (or noise) in the abundance of transcription factors within network motifs is architecture-dependent and sharply contrasts with its buffering effect on noise in the expression of isolated genes. Overall, our results show that biomolecular phase separation acts as a physical principle for tuning stability and noise in gene regulatory networks, providing new insights into cellular decision-making.

## I. INTRODUCTION

Single cells make diverse functional decisions using gene regulatory networks (GRNs). GRNs consist of molecular regulators such as transcription factors (TFs) that interact with each other to control gene expression [1–3]. Certain recurring patterns called motifs are overrepresented in GRNs [2, 4]. Canonical motifs include autoregulation and feed-forward loops, which tune response times, stabilize expression, or filter noise [2, 4–6]. In synthetic biology, the toggle switch and the repressilator motifs have been engineered to demonstrate how simple regulatory architectures can generate bistability and oscillations [6–9]. These motifs have been harnessed as modular building blocks for designing programmable circuits for biosensing and therapeutic control, and they have also been instrumental in unraveling the governing principles of cellular decision-making [10, 11]. Analogous motifs exhibiting bistable or oscillatory dynamics are indeed found in natural systems. Switch-like (bistable) motifs play central roles in regulating processes such as the cell cycle in yeast [12], the lysis–lysogeny switch in bacteriophage *λ* [13, 14], the lac operon in Escherichia coli [15, 16], and various cellular signaling pathways [17–20]. Oscillatory motifs, on the other hand, underlie rhythmic behaviors including circadian cycles, stress responses, and pulsatile signaling in both prokaryotic and eukaryotic systems [21–26].

Cells regulate the dynamical behavior of gene regulatory motifs through multiple processes, including transcriptional, post-transcriptional, translational, post-translational, and TF degradation mechanisms, as well as through sequestration and subcellular localization [2, 27–31]. Global resource constraints, such as the limited availability of RNA polymerases, ribosomes, and ATPs, also shape gene expression dynamics [32–35]. Among these regulatory strategies, controlling TF availability in cell is particularly central to gene regulation. For example, binding of TFs to high-affinity, non-functional genomic decoy sites sequesters TFs and reduce the concentration of freely available TFs for a target promoter [36–38]. Tuning the number of decoy sites and the affinity of transcription factors (TFs) for these decoys leads to significant changes in the dynamics of gene regulatory motifs; for instance, TF sequestration by decoys can shift basin boundaries and modify the stability of fixed points in a toggle switch [39].

In recent years, phase separation of TFs has emerged as an alternative mechanism for regulating TF concentration in cells (Fig. 1a). Above a critical concentration (also known as the saturation concentration), TFs can undergo phase separation, leading to the formation of a dense phase that coexists with a surrounding dilute phase. This mechanism spatially compartmentalizes regulatory components and thereby influences gene expression [40, 41]. A growing body of work has shown that phase-separated condensates of TFs play scores of regulatory roles in transcription. For instance, activator condensates, often formed at super-enhancers near promoters, enhance gene expression by locally enriching RNA polymerase II and coactivators [42–44]. Moreover, phase separation of TFs has been linked to transcriptional memory, bistability, and noise buffering, suggesting that condensates can stabilize gene expression states and enhance regulatory robustness [45–48]. Another study has shown that phase separation can suppress transcriptional bursting and decrease cell-to-cell variability in gene expression [49]. Phase separation can also drive phenotype switching in autoregulatory systems by coupling condensate dynamics to regulatory feedback [50]. Despite these advances, most existing studies have focused on single-gene systems or specific transcriptional events [45–47, 49, 50], leaving unresolved how phase separation influences the dynamics of gene regulatory motifs in cells.

**FIG. 1.**
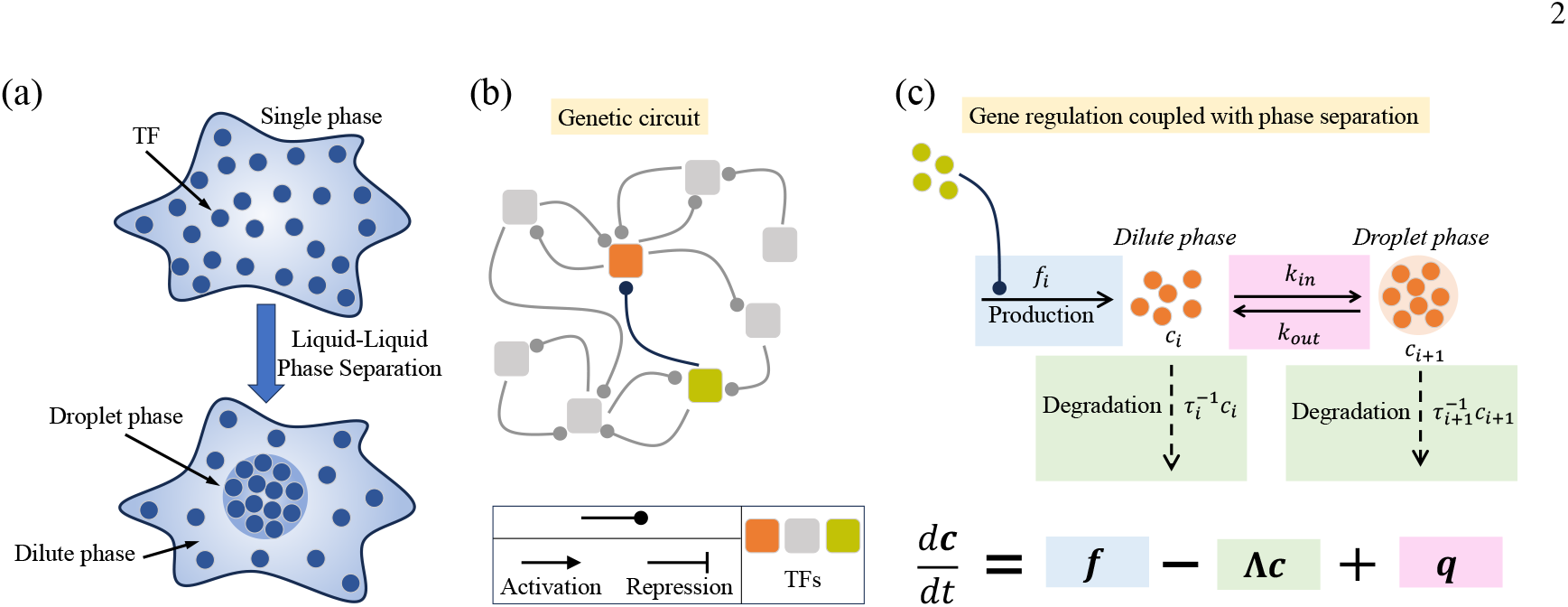
Gene regulation coupled with phase separation. (a) Schematic of a cell in which TFs undergo phase separation into dilute and droplet phases. (b) Representation of a generic gene regulatory circuit composed of interacting TFs. (c) Illustration of TF production and degradation processes coupled to phase separation. The color blocks correspond to the respective terms in Eq. (2).

The goal of this manuscript is to develop a theoretical framework that enables the study of how protein phase separation influences the dynamical behavior of GRNs (Figs. 1b,c). We apply this framework to study two well-known motifs, the toggle switch and the repressilator (Fig. 2). The toggle switch, composed of two mutually repressing genes, underlies binary cell-fate decisions and serves as a model for memory and decision-making in synthetic biology [7]. The repressilator, a cyclic three-gene network, generates sustained oscillations and represents the minimal architecture for rhythmic gene expression [8]. To elucidate how phase separation of the TFs associated with these motifs modifies their dynamics, we examine two scenarios: (i) single-TF phase separation and (ii) multi-TF phase separation. Using mean-field theory, we show that phase-separating TFs reshape the dynamical behavior. In particular, when multiple TFs undergo phase separation, the one with the lowest saturation concentration dominates the overall system dynamics. Stochastic simulations further uncover how phase separation tunes noise in TF abundance. A phase-separating TF in the toggle switch primarily modulates fluctuations of its own high-expression state and the low-expression state of its repressor. In addition, the phase-separating TF in the repressilator reduces its own amplitude fluctuations while amplifying those of the other TFs in the circuit. These results show that the effect of phase separation on gene expression noise is dictated by the architecture of gene regulatory networks. Overall, our theoretical framework offers a means to analyze and control the dynamical behavior of gene regulatory networks when their constituent molecular components undergo phase separation within cells.

**FIG. 2.**
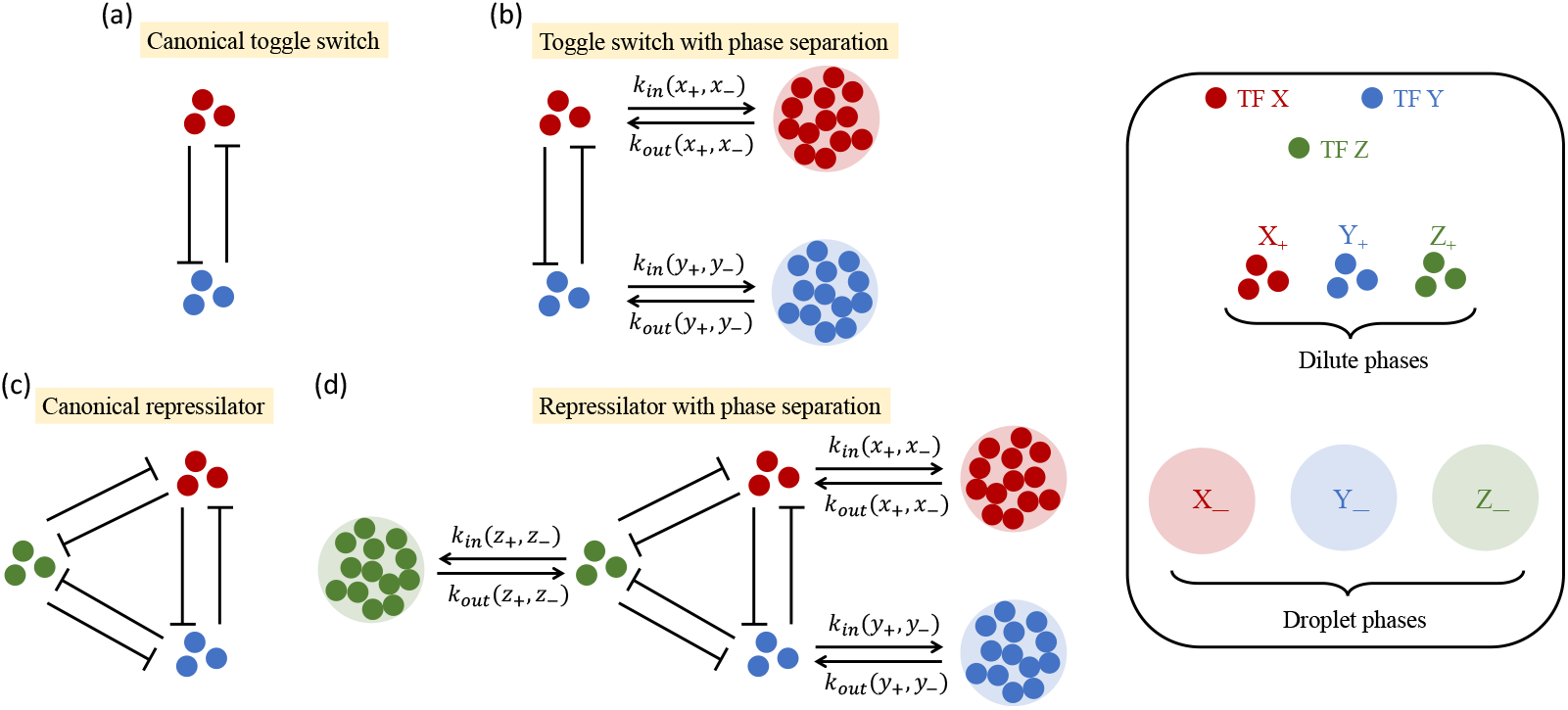
Schematics of regulatory motifs coupled with phase separation. (a) Canonical toggle switch consisting of two TFs, X and Y, that mutually repress each other. (b) Toggle switch with phase separation. In this case, one TF may undergo phase separation while the other remains in the well-mixed state, or both TFs may phase separate simultaneously. (c) Canonical repressilator where the three prtoeins, X, Y, and Z, mutually represss each other. (d) Repressilator with phase separation, illustrated with all three TFs undergoing phase separation. In the analysis, however, the different scenarios of one, two, or three TFs undergoing phase separation are considered separately.

## II. A GENERAL MODEL OF BINARY PHASE SEPARATION COUPLED TO GENE REGULATORY NETWORKS

To investigate how phase separation of TFs influences the dynamics of gene regulatory networks, we construct a theoretical framework that integrates a minimal model of binary phase separation with a GRN of arbitrary complexity (Figs. 1a-c). Our formulation begins with a simple model of binary phase separation that describes partitioning between dilute and droplet phases under the constraint of constant total TF number (Fig. 1a). We note that we adopt this binary mixture model following the formulation of Klosin et al. [47]. A complete thermodynamic description of binary phase separation and the corresponding kinetics of molecular exchange is provided in the *SI Text*. We develop a framework by coupling this model of binary phase separation to an arbitrarily complex GRN and apply it to dissect the dynamical behavior of toggle switch and repressilator. The general GRN (Fig. 1b) is composed of *m* TFs such that ***c*** = (*c*_1_, *· · ·, c*_*m*_)^⊤^ represent their copy number vector (state space). The mean-field dynamics of the gene regulatory network can be written as,

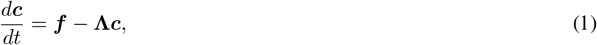

where, ***f*** = (*f*_1_, *· · ·, f*_*m*_)^⊤^ consists of the production rates of different TFs and 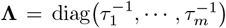 is the diagonal matrix of first-order degradation rate constants. To incorporate phase separation, we expand the state space by representing each phase-separated TF with two adjacent variables – the dilute phase copy number followed by the droplet phase copy number. With this convention, the enlarged state space is ***c*** = (*c*_1_, *· · ·, c*_*n*_)^⊤^ with *n* ≥ *m*. While there remain *m* distinct TFs, phase separation increases the dimension of the state space. In particular, equality (*n* = *m*) holds when no TF phase separates, whereas strict inequality (*n > m*) arises when one or more TFs undergo phase separation. Now, for every phase-separating TF, there exists an index *i* in the state space such that (*c*_*i*_, *c*_*i*+1_) represents its (dilute, droplet) pair. Additionally, the regulatory interactions in the production terms *f*_*i*_ use only dilute phase copy numbers as the molecules inside the droplets do not participate in transcription. Considering all these aspects, the deterministic dynamics with phase separation can now be written as (Fig. 1c),

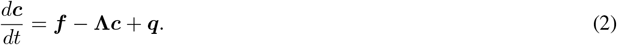

where ***f*** = (*f*_1_, *· · ·, f*_*n*_)^⊤^ and 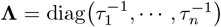. Note that we consider non-zero degradation rates for the TFs in the droplet phase. In the above equation, the vector ***q*** = (*q*_*c*_, *· · ·, q*_*c*_)^⊤^ accounts for the net molecular flux due to exchange of TF molecules between dilute and droplet phases and is defined by (Fig. 1b),

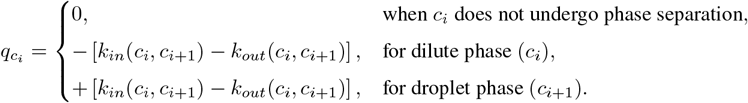

Here, *k*_*in*_ and *k*_*out*_ refer to the kinetic rates of TF exchange from the dilute to droplet phase and vice versa, respectively. These rates depend on the instantaneous copy numbers of the dilute and droplet phases, which depend on the dynamics of gene regulation (Fig. 1b). However, their explicit forms are defined in terms of the thermodynamic and kinetic parameters of the binary phase separation model with constant TF number (see *SI text*). In that formulation, the onset of phase separation is characterized by the critical TF copy number (*p*^∗^). When this framework is coupled to gene regulation, TF copy numbers are no longer fixed, and we denote the corresponding threshold for the *i*-th TF in the circuit by 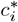. Therefore, *p*^∗^ defines the global onset threshold in the simplified binary mixture, while 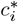 serves as its TF-specific analogue in the regulatory circuits.

To incorporate stochasticity, we formulate the corresponding chemical master equation (CME) for the joint probability distribution 𝒫 (***c***, *t*). We now define the index set 𝒥 = { *i* | (*c*_*i*_, *c*_*i*+1_) forms a dilute-droplet pair of the same TF }, which collects all positions in ***c*** where *c*_*i*_ denotes the dilute phase copy number and *c*_*i*+1_ represents the corresponding droplet phase copy number of the same TF. For example, consider a genetic circuit with three TFs X, Y, and Z (*m* = 3), where Y and Z undergo phase separation (i.e., *n* = 5). The state vector becomes ***c*** = (*c*_1_, *c*_2_, *c*_3_, *c*_4_, *c*_5_)^⊤^, with *c*_1_ denoting the copy number of X, (*c*_2_, *c*_3_) representing Y in dilute and droplet phases, and (*c*_4_, *c*_5_) representing Z in dilute and droplet phases. In this case, the set of dilute–droplet indices is 𝒥 = {2, 4}, corresponding to the dilute phases of Y and Z. The CME is given by,

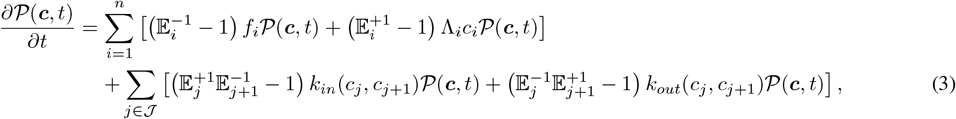

where 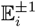 denote the step operator defined as, 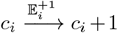 and 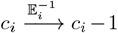. In Eq. (3), the first sum accounts for production and degradation of all species, and the second sum includes only phase-exchange events between dilute-droplet pairs 𝒥. This stochastic description explicitly accounts for intrinsic noise arising from finite copy numbers and discrete reaction events. We now apply this general framework to specific cases – the toggle switch and repressilator.

### A. Toggle switch

A toggle switch consists of two TFs, X and Y, that mutually repress each other”s expression (Fig. 2a). The mean-field dynamics of this system can be described by Eq. (1), where,

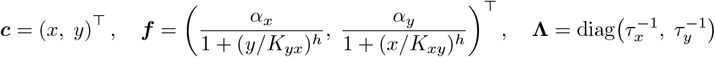

Here, *x* and *y* denote the copy numbers of TFs X and Y, respectively. We note that in the general formulation, we use *c*_*i*_ to represent the copy number of the *i*-th TF, but in specific motifs, we adopt simpler lowercase symbols for readability. Here, the parameters *α*_*x*_ and *α*_*y*_ represent their maximal production rates. The terms *K*··· denote the dissociation constant of a TF regulating its target gene. For example, *K*_*yx*_ refers to the dissociation constant of TF Y regulating the gene X, and *K*_*xy*_ can be defined analogously. Here, *h* is the Hill coefficient capturing cooperativity in TF binding to DNA.

To investigate the effect of phase separation on the dynamics of toggle switch, we incorporate the effect of phase separation of TFs into the mean-field dynamics of toggle switch under two scenarios: (i) only TF Y undergoes phase separation, and (ii) both X and Y undergo phase separation (Fig. 2b). In each case, gene regulation is mediated by TFs in the dilute phase, since molecules partitioned into the droplet phase are spatially sequestered from promoter regions and remain transcriptionally inactive. The generalized mean-field dynamics for these two scenarios are described by Eq. (2). To be precise, for case (i), the state vector becomes ***c*** = (*x, y*_+_, *y*_−_)^⊤^, with *y*_+_ and *y*_−_ being the population of TF Y in the dilute and droplet phases, respectively. The other terms are

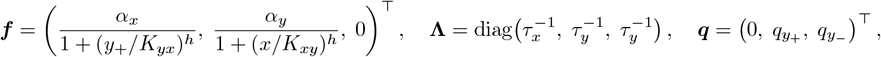

where 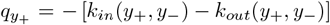 and *q*_*y−*_ = *k*_*in*_(*y*_+_, *y*_*−*_) *−k*_*out*_(*y*_+_, *y*_*−*_). We note here that the degradation rate of TFs in droplet phase is assumed to be equal to that in the dilute phase. The onset of phase separation for TF Y is specified by its threshold copy number *y*^∗^, which is the motif-specific analogue of the general threshold 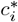 defined earlier. However, for case (ii), the terms of Eq. (2) are given by,

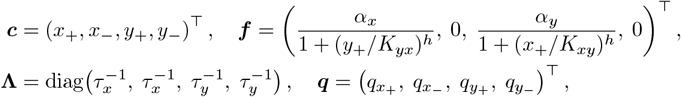

where 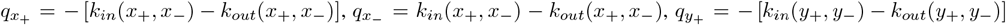, and 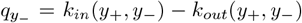 Similar to the former case, we use equal degradation rates for TFs in both the dilute and droplet phases. The phase separation thresholds for TFs X and Y are denoted by *x*^∗^ and *y*^∗^, respectively.

The stochastic dynamics in both scenarios are governed by the generalized CME, Eq. (3). We simulate the dynamics using Gillespie”s stochastic simulation algorithm (SSA) [51, 52]. The propensities for the reaction channels for each model are given in *SI text*. The specific parameter values used are also summarized in the *SI text*.

### B. Repressilator

A repressilator consists of three TFs – X, Y, and Z – where each TF represses the expression of the next in a cyclic fashion (Fig. 2C). The deterministic dynamics is governed by Eq. (1), where the following definitions hold,

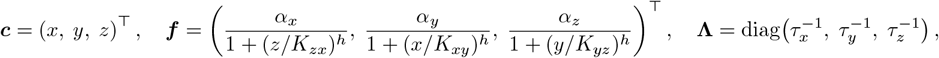

where *x, y*, and *z* represent the TF copy number of X, Y, and Z, respectively. The parameters 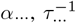 and *K*… carry the analogous interpretation as described for toggle switch. In this system, we introduce three scenarios: (i) only Z phase separates, (ii) Y and Z phase separate, and (iii) all the three TFs phase separate. The mean-field dynamical equation is given by Eq. (2) for the three given conditions. The matrices involved for these three cases are given as follows: In model (i),

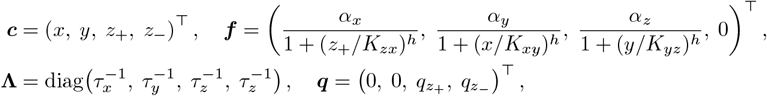

where, 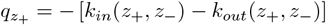 and 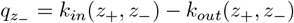. In model (ii),

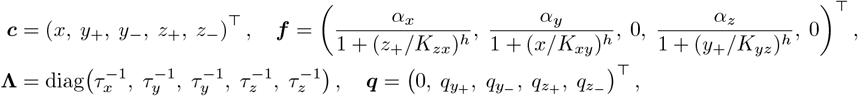

where, 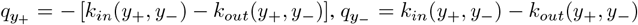, and 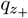 and 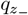 follow from (i). In model (iii),

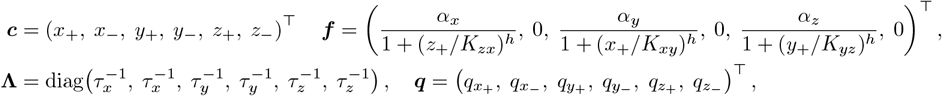

where, 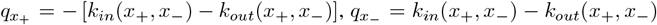, and 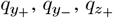 and 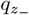 follows from (ii). We denote the critical thresholds of phase separation for these cases as follows: *z*^∗^ for (i), (*y*^∗^, *z*^∗^) for (ii), and (*x*^∗^, *y*^∗^, *z*^∗^) for (iii). The equation governing stochastic dynamics of TFs for these cases is given by Eq. (3). We simulate their dynamics using SSA [51, 52]. The propensities for the reaction channels corresponding to the different processes for each model are given in *SI text*.

Using mean-field analysis and stochastic simulations, we probe the impact of phase separation of TFs on the dynamical behavior of toggle switch and repressilator, as discuss in the ensuing sections.

## III. RESULTS

### A. Reshaping the dynamical landscape of TFs due to phase separation

#### Toggle switch

We begin by analyzing the mean-field dynamics of the toggle switch (Fig. 2a). The dynamical landscape for the canonical toggle switch is obtained from Eq. (1), while the landscapes for the phase-separating cases are generated using Eq. (2). The canonical toggle switch exhibits two stable fixed points (attractors) and an unstable fixed point (see Fig. 3a). At the stable states, the TFs X and Y adopt mutually exclusive expression patterns: {*x=high, y=low*} and {*x=low, y=high*}. At the unstable point, the TFs are expressed equally at some intermediate levels. Each stable state is associated with a basin of attraction, defined as the region of concentration space from which trajectories converge to that state. The two basins corresponding to {*x=high, y=low*} and {*x=low, y=high*} are separated by a linear separatrix and hence are symmetric (Fig. 3a). We now investigate how introducing phase separation in one and both TFs perturbs this inherent symmetry in the dynamical landscape of the toggle switch.

**FIG. 3.**
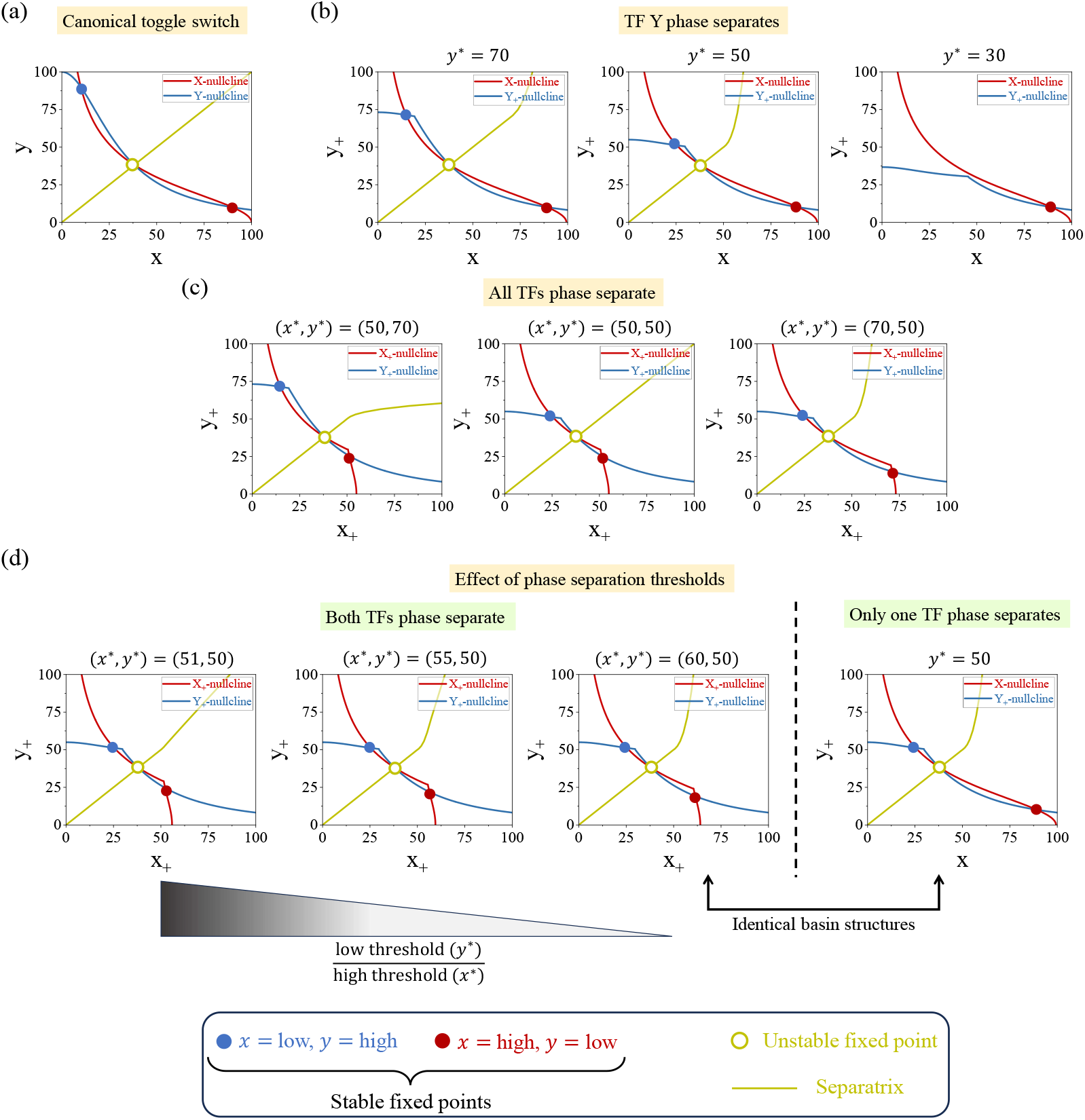
Phase separation reshapes the dynamical landscape of the toggle switch. (a) Canonical toggle switch with symmetric basins separated by a linear separatrix. (b) Toggle switch with phase separation in TF Y. (c) Toggle switch with phase separation in both TFs X and Y. (d) Toggle switch showing the effect of unequal thresholds, where the lower threshold dominates the dynamics. Panel a is generated using Eq. (1), whereas panels b–d are generated using Eq. (2). Parameter values are provided in the *SI text*.

First, we allow TF Y to undergo phase separation at different thresholds *y*^∗^ (Fig. 3b). As *y*^∗^ decreases, the fixed point {*x=low, y=high*} shifts downward along the X-nullcline, whereas the other fixed point {*x=high, y=low*} remains unchanged. This shows that phase separation of Y selectively perturbs its own high-expression state and the low-expression state of its repressor. At the same time, the separatrix bends toward the {*x=low, y=high*} basin, with the curvature becoming more pronounced at lower thresholds. This bending shrinks the basin of attraction of{*x=low, y=high*} state, while expanding the other basin. A smaller basin implies that fewer initial conditions lead to convergence to the corresponding attractor, thereby biasing the dynamics toward the opposite state. At *y*^∗^ = 30, the threshold is close to the unstable point, which causes {*x=low, y=high*} state to disappear and the toggle switch becomes monostable. Phase separation of TF Y, thus, breaks the inherent symmetry of the toggle switch and enhances the stability and dominance of the {*x=high, y=low*} state.

When both X and Y undergo phase separation, the thresholds *x*^∗^ and *y*^∗^ reshape the dynamical landscape of the toggle switch (Fig. 3c). When the thresholds are unequal, the lower one dominates, and the separatrix bends towards the basin where the corresponding TF – the one having lower threshold – is highly expressed. When the thresholds are equal (*x*^∗^ = *y*^∗^), the landscape regains symmetry, but the stable points are displaced compared to the non-phase-separating toggle switch. Thus, unlike the single-TF case, where phase separation only perturbs the high-expression state of the TF involved, the dual-TF case is governed by threshold asymmetry, with the lower threshold remodeling the orientation of the separatrix and the basin geometry.

A key result emerges when the thresholds are well-separated. In this regime, the TF with lower phase separation threshold governs the dynamical landscape of the system. As shown in Fig. 3d, when the ratio of the low to the high threshold is very small, the resulting basins of attraction become indistinguishable from the basin structure in the single-TF case where only the lower threshold is present. Although the positions of the stable points are not identical, the geometry of the basins of attraction is governed by the TF with lower threshold. This reveals an interesting dynamical signature of phase separation: when thresholds differ substantially, the lower threshold dominates the dynamics and determines the effective switching behavior of the toggle switch.

#### Repressilator

We next analyze the mean-field behavior of the repressilator in both the absence and presence of phase separation by generating time series and corresponding phase-space trajectories. For a canonical repressilator, we use Eq. (1), while the phase-separated cases are modeled with Eq. (2). In these phase-separated cases, the dynamical behavior is determined solely by the dilute-phase TF populations, because molecules sequestered in droplets are transcriptionally inactive. In the absence of motif phase separation, the three TFs X, Y, and Z generate sustained oscillations with equal amplitude and frequency for when the biophysical parameters characterizing the models are the same, yielding a symmetric limit cycle (Fig. 4). Note that repressilator exhibits stable limit cycles owing to a supercritical Hopf bifurcation of the stable equilibrium point. When phase separation is introduced, however, the symmetry of the limit cycle is broken (Fig. 4a–c). The resulting asymmetry manifests as changes in oscillation amplitude and period along the directions of the phase-separating TFs. For instance, when only TF Z undergoes phase separation, the distortion originates specifically from Z (Fig. 4a). Comparable asymmetries arise when two TFs, or all three, phase separate (Fig. 4b,c).

**FIG. 4.**
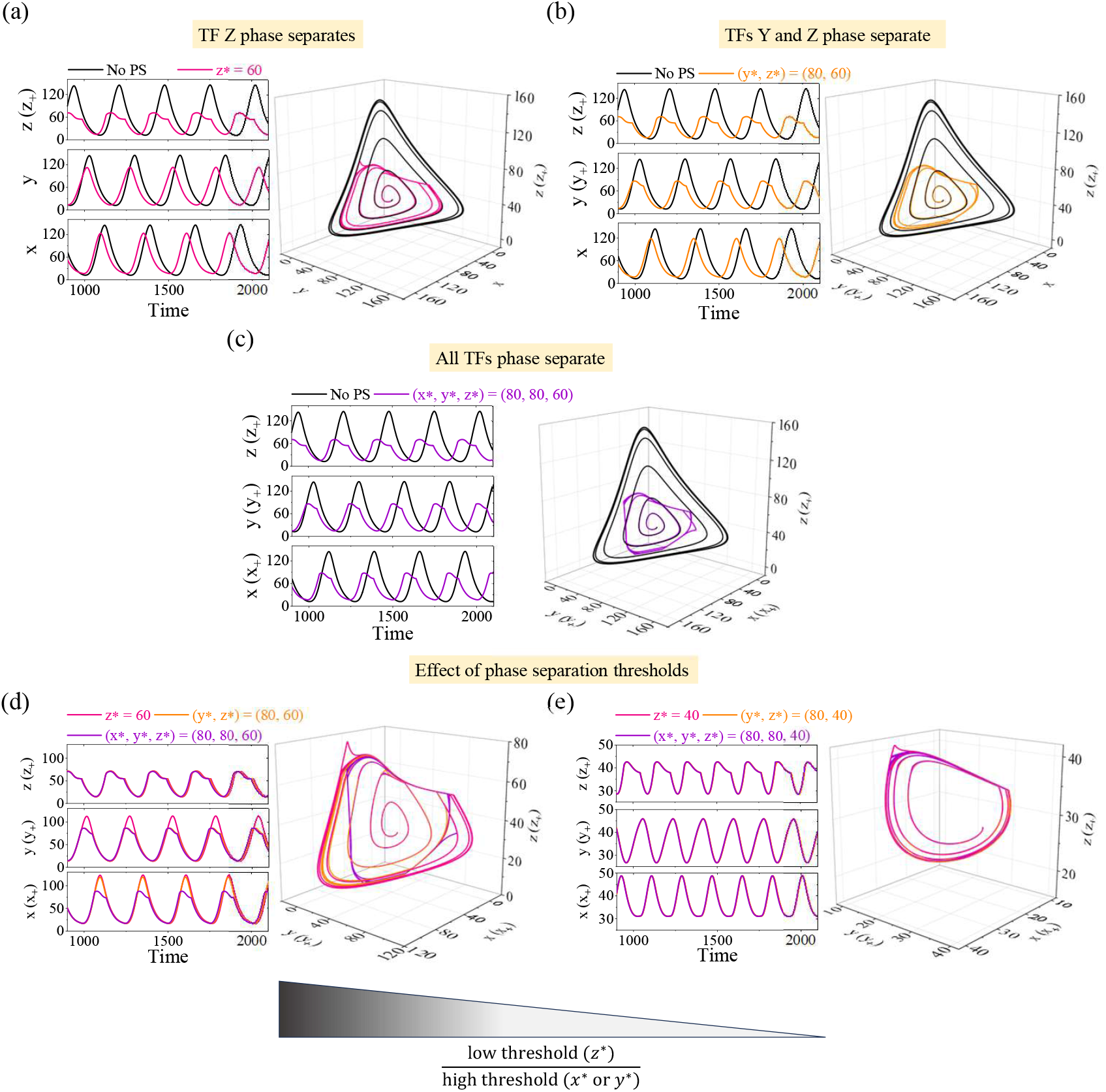
Phase separation reshapes the dynamical landscape of the repressilator. In panels a-c, the canonical repressilator is compared with three different scenarios of phase separation. (a) Only TF Z phase separates, (b) TF Y and Z phase separate, and (c) All TFs phase separate. (d) Compares the three scenarios of phase separation with moderate differences in phase separation thresholds. (e) Compares the same scenarios with strong differences in phase separation thresholds. The abbreviation “No PS” denotes the canonical repressilator without phase separation. The No PS result is obtained from Eq. (1), whereas the results for the phase-separated repressilators are obtained from Eq. (2). Parameter values are provided in the *SI text*.

A distinct feature emerges when comparing the three scenarios of phase separation – one TF, two TFs, and all three TFs within the same frame (Fig. 4d,e). We find that in Fig. 4d, where the thresholds differ only moderately (a moderate ratio of low to high threshold), the oscillations deviate from the symmetric limit cycle but do not collapse onto the behavior of any single-TF case. In contrast, Fig. 4e illustrates the regime of strong threshold separation (small ratio of low to high threshold), where the oscillations are dictated almost entirely by the TF with the lowest threshold. In this regime, the amplitude and period exactly match those of the corresponding single-TF case, and the influence of higher thresholds becomes negligible.

These observations highlight a broader principle linking the toggle switch and repressilator. In a toggle switch, when both transcription factors undergo phase separation, the one with the lower saturation concentration predominantly governs the system”s dynamics. The TF with higher saturation concentration does not affect the basin geometry. Similarly, in the case of a repressilator, when the saturation concentrations are well separated, the transcription factor with the lowest saturation concentration determines both the amplitude and period of oscillations, while those with higher saturation concentrations have negligible influence on the oscillatory dynamics. Thus, despite the structural differences between the gene regulatory networks we consider, phase separation of TFs imposes a common regulatory rule: when saturation concentration for bulk phase separation of multiple TFs in a regulatory motif are well separated, TF with the lowest saturation concentration effectively governs the dynamics of the system.

### B. Regulation of noise in TF abundance by phase separation

#### Toggle switch

We next examine how phase separation of TFs impact fluctuations (or noise) in the TF abundance for the toggle switch. To assess noise around each stable fixed point, we generate stochastic time series of TFs X and Y using SSA (Fig. 5a). For phase-separated systems, we track only the dilute phase copy numbers. From the time series, we extract the corresponding steady-state distributions. Because the toggle switch is bistable, noise is evaluated separately for the high- and low-expression states of each TF (Fig. 5a). Noise is, then, quantified using the coefficient of variation (CV), defined as the ratio of the standard deviation to the mean for each expression state. This approach enables systematic comparison of molecular noise across conditions with and without phase separation and for different thresholds.

**FIG. 5.**
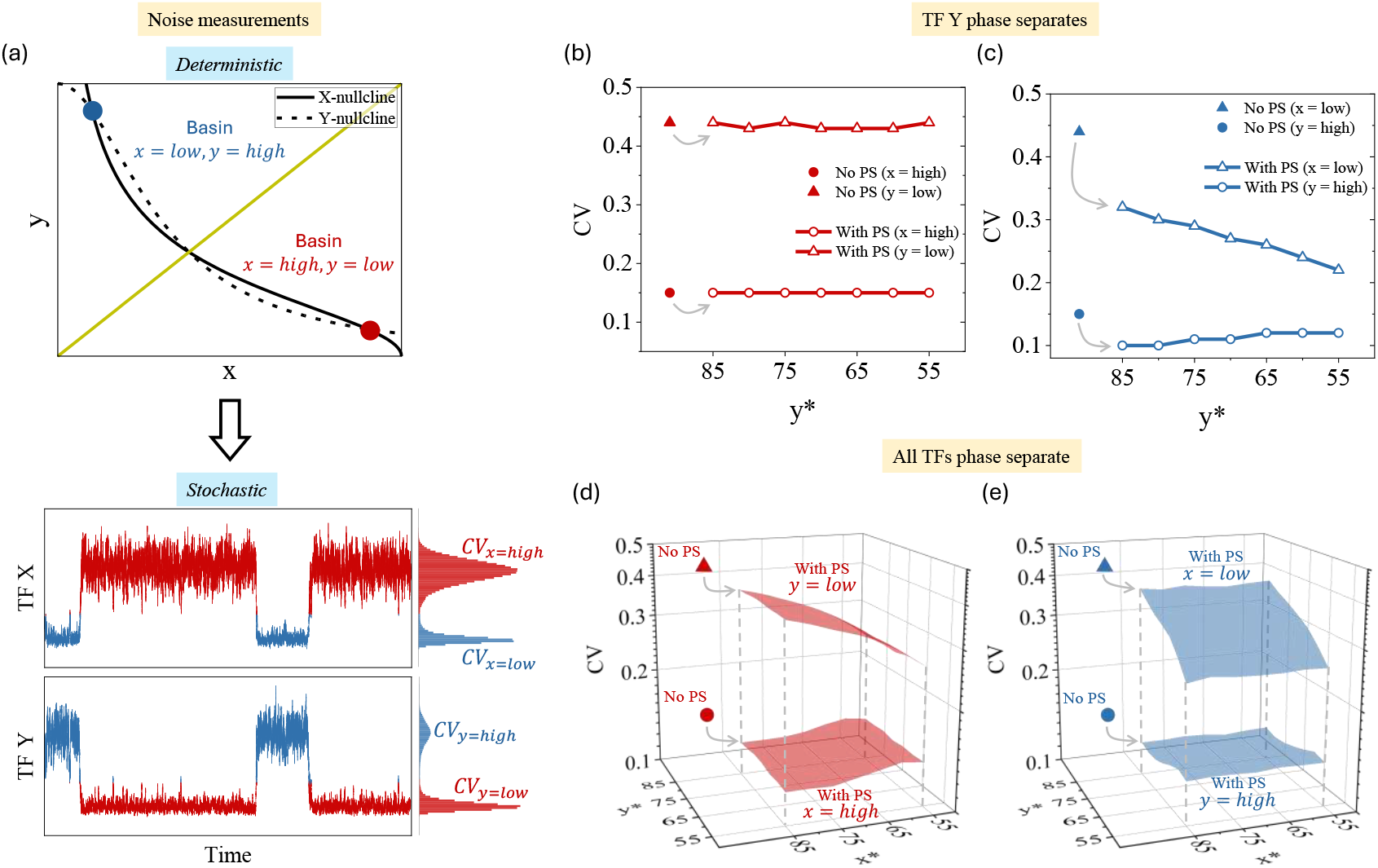
Effects of phase separation on the noise properties of toggle switch. (a) Schematic of the noise measurement procedure. Stochastic time series are generated for each stable fixed point, and the corresponding distributions are used to compute the CV (standard deviation / mean) as a measure of molecular noise. (b-c) Effect of phase separation in TF Y on noise at different thresholds. Panel b shows the CV for the state {*x=high, y=low*}, while panel c shows the CV for the state {*x=low, y=high*}. (d-e) Effect of phase separation in both TFs X and Y on noise across different thresholds. Panel d quantifies the CV for the state {*x=high, y=low*}, and panel e quantifies the CV for the state {*x=low, y=high*}. In panels b-e, “No PS” denotes the canonical toggle switch without phase separation. Reaction propensities and parameter values are provided in the *SI text*.

We first consider the case when only TF Y undergoes phase separation. By varying the threshold *y*^∗^, we compare the noise profiles with those of the canonical toggle switch in absence of phase separation. We find that the noise associated with the *x=high* and *y=low* states remain essentially unchanged across all values of *y*^∗^ and are comparable to the corresponding states of the canonical toggle switch (Fig. 5b). This invariance arises because the fixed point {*x=high, y=low*} does not change when Y undergoes phase separation at different thresholds (see Fig. 3b). In contrast, noise in both *x=low* and *y=high* states are reduced relative to the canonical toggle in absence of phase separation. This reduction stems from sequestration of Y into the droplet phase, which buffers fluctuations in its dilute phase population. Moreover, the effect of *y*^∗^ differs between the two states: the noise in the *x=low* state decreases substantially with decreasing *y*^∗^, while the noise in the *y=high* state increases slightly. The observations are consistent with the deterministic shift of the fixed point {*x=low, y=high*} along the X-nullcline as *y*^∗^ is lowered (see Fig. 3b). These findings reveal a key feature of the toggle switch: a phase-separating TF primarily regulates the noise of its own high-expression state and also influences the low-expression state of its repressor.

Next, we consider the case where both TFs undergo phase separation. We observe that the noise associated with *x=high* and *y=low* states are lower compared to their corresponding counterpart for the canonical toggle switch (Fig. 5d). Moreover, the noise levels are selectively regulated by the phase separation threshold of TF X. Here, the noise of *y=low* is reduced significantly while the same for *x=high* is increased slightly as *x*^∗^ decreases (Fig. 5d). A complementary pattern is observed for the noise of *x=low* and *y=high* states, where the threshold *y*^∗^ primarily controls the noise levels (Fig. 5e).

The emergent picture for toggle switch is that phase separation consistently reduces the overall noise of TFs in the dilute phase. The detailed effect on individual TFs, however, is asymmetric, where a phase-separating TF primarily increases the noise of its own high-expression state and reduces the noise of the low-expression state of its repressor.

#### Repressilator

Next, we analyze how phase separation influences fluctuations in the biophysical properties of oscillatory cycles generated by the repressilator. Stochastic time series of TFs X, Y, and Z are gerenated using SSA for both canonical repressilator (Fig. 6a) and repressilator where one or more TFs undergo phase separation. For phase-separated systems, only the dilute phase TF abundances are tracked. In each time series, individual oscillation cycles are identified to compute the amplitude and period distributions. The distributions for the canonical repressilator are shown in Fig. 6a, and the same procedure is applied to the phase-separating cases. The details of amplitude and period measurements from the stochastic time series are provided in the *SI text*. From these distributions, we calculate the CV for amplitude and period to quantify the relative fluctuations between oscillation cycles.

**FIG. 6.**
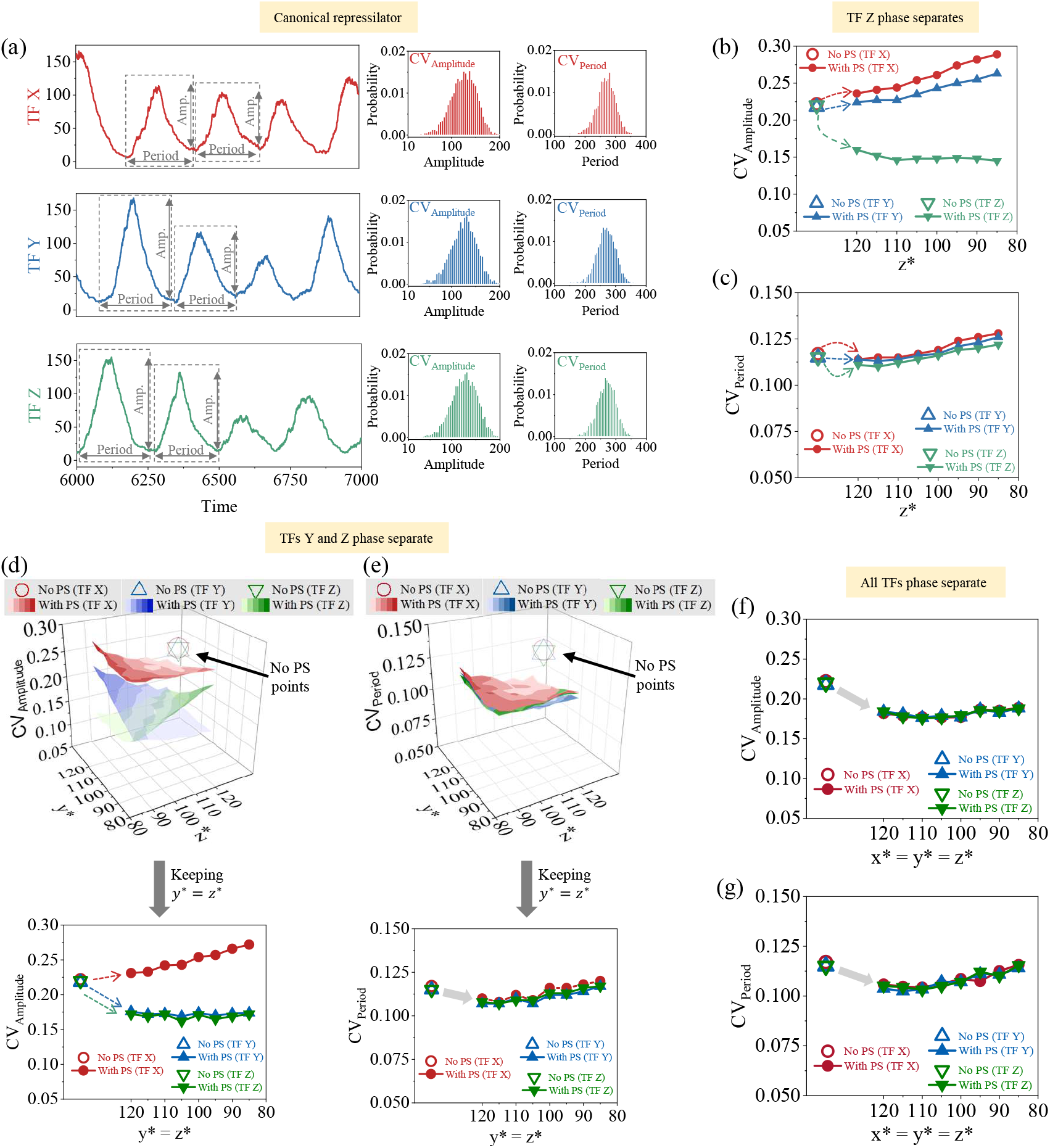
Effects of phase separation on the noise properties of repressilator. (a) Schematic of the noise measurement procedure. Stochastic time series are generated for each TF, and the corresponding amplitude and period distributions are used to compute the CV (standard deviation divided by mean) as a measure of fluctuations in the oscillation. (b-c) Effect of phase separation in TF Z on amplitude and period noise at different thresholds. (d-e) Effect of phase separation in TFs Y and Z on amplitude and period noise across different thresholds. (f-g) Effect of phase separation in all TFs on amplitude and period noise across different thresholds. In panels b-g, “No PS” denotes the canonical repressilator without phase separation. Reaction propensities and parameter values are provided in the *SI text*.

We first consider the case where TF Z undergoes phase separation. We find that the amplitude noise of Z decreases upon phase separation, whereas that of X and Y increases relative to the canonical repressilator. These effects become more pronounced as the threshold *z*^∗^ decreases, indicating that the more readily Z is sequestered into the droplet phase, the lower its amplitude noise, and the higher the amplitude noise of X and Y (Fig. 6b). However, the noise in period of all TFs remains largely unchanged with phase separation (Fig. 6c). These behaviors can be explained by the buffering effect of phase separation and the architecture of the repressilator. When Z phase separates, molecular abundance of Z in the dilute phase becomes buffered, leading to a reduction in its amplitude noise. The reduced level of Z in the dilute phase, in turn, weakens its repression strength on X, making X more susceptible to intrinsic fluctuations, which then propagate to Y through the cyclic repression loop. Consequently, amplitude noise increases for X and Y. Despite these changes in amplitude, the noise in period remains nearly constant because phase separation tunes the effective repression strengths without altering the intrinsic synthesis and degradation rates that set the oscillation timescale.

When both Y and Z undergo phase separation, the repressilator exhibits trends similar to the case of single-TF phase separation. We find that the amplitude noise of Y and Z are reduced relative to the canonical repressilator, reflecting the buffering effect of their dilute-phase abundances. However, the amplitude noise of X increases as the thresholds *y*^∗^ and *z*^∗^ decrease, owing to the weakening of repression strengths of Y and Z (Fig. 6d). The period noise for all TFs remains largely unaffected (Fig. 6e), consistent with the fixed synthesis and degradation rates that determine the oscillation timescale. These patterns persist when all three TFs phase separate, where amplitude noise is reduced for all TFs compared with the canonical repressilator, while period noise remains nearly unchanged (Figs. 6f,g).

To summarize, our results for the repressilator show that phase separation selectively modulates amplitude fluctuations of TFs in the feedback loop. A phase-separating TF primarily suppresses its own amplitude noise while amplifying that of the other TFs through regulatory coupling. When multiple TFs undergo phase separation, these effects combine to produce a collective buffering of fluctuations, leading to a global reduction in amplitude noise across the network.

One of the key findings of our study is that the effect of phase separation on expression noise of genes associated with a regulatory network depends strongly on the network architecture. This behavior is sharply distinct from how phase separation buffers noise in expression of isolated genes.

## IV. DISCUSSION

In this work, we develop a theoretical framework that couples gene regulatory networks with binary phase separation and apply it to study the dynamical properties of canonical network motifs, namely the toggle switch and repressilator. We show that phase separation of TFs reshapes both the dynamical landscape and the noise properties of these motifs. The key determinant of motif dynamics is the relative values of phase separation thresholds (critical concentrations for the onset of phase separation) of multiple TFs. When the thresholds are well-separated, the TF with the lowest threshold dominates the system dynamics, governing basin geometry in the toggle switch and controlling oscillation amplitude and period in the repressilator. We also find that hase separation of TFs consistently buffers molecular fluctuations in the abundance of TFs in dilute phase. However, motif architecture dictates how the fluctuations propagate across the network. For instance, in toggle switch, a phase-separating TF differentially regulates the noise in the high-expression state of its own and the low-expression state of the opposing TF. In the repressilator, a phase-separating TF suppresses the amplitude noise of its own while amplifying the same of the other TFs in the motif. This finding demonstrates that the way phase separation impacts noise among the TFs associated with a gene regulatory network is sharply distinct from its buffering effect on expression noise for isolated genes. Only when all TFs phase-separate does the amplitude noise of each get suppressed. Together, these results establish that phase separation of TFs shape both the dynamical behaviour and noise in TF abundance, providing a tuning knob to control the dynamics of gene regulatory networks.

Our study builds on previous findings on the role of phase separation in transcriptional regulation and advances them by linking molecular-scale phenomena to network-level behavior. Previous studies demonstrated that TFs and coactivators can form condensates that modulate gene expression, either by concentrating factors at promoters or by buffering fluctuations in their availability [45, 47, 53]. However, most existing analyses have focused on individual genes [47, 49, 50]. Our theoretical framework connects binary phase separation with canonical gene regulatory motifs, showing how phase separation reshapes network dynamics and fluctuations in TF abundance. This framework, thus, provides a systematic way to investigate how phase separation-mediated compartmentalization of TFs can regulate variability in gene expression depending on network architecture.

A recent theoretical study [39] examined how TF sequestration by genomic decoy sites affects the dynamics of toggle switch and repressilator. The study showed that TF binding to decoy sites reshapes the basin of attraction in the toggle switch and modify oscillatory behavior of the repressilator by controlling the pool of free TFs. Lyons et al. [9] similarly showed that coupling a toggle switch to a downstream load sequesters regulatory molecules, skewing its potential landscape and modifying bistability. Interestingly, these studies highlight that molecular sequestration of the different TFs associated with the gene regulatory network, whether through decoys or downstream interactions, collectively reshape the dynamical behavior of regulatory motifs. In sharp contrast, when all the TFs undergo phase separation, the dynamics of toggle switch and repressilator are dictated by the TF with the lowest threshold concentration for phase separation. Furthermore, our stochastic analysis quantifies how phase separation influences the noise characteristics of these motifs. We also analyze, in SI Text, how the interplay between the timescales of TF degradation and their diffusion-limited phase separation affects the dynamical behavior of the toggle switch and repressilator. When the diffusion is much faster than TF degradation, the basin structure of the toggle switch becomes asymmetric, and the oscillation amplitude in the repressilator is effectively buffered.

In synthetic biology, phase separation offers a new layer of tunability to modulate gene regulatory dynamics. Unlike traditional regulatory parameters such as promoter strength or degradation rate [7, 8, 54], phase separation introduces threshold-like non-linearities that can be achieved by appropriately designing synthetic proteins. Our results suggest that by tuning phase separation thresholds, one could bias cell-fate switches and adjust oscillatory precision in synthetic systems. For instance, engineering a toggle switch with phase-separating TFs could stabilize the attractors against fluctuations. Similarly, designing repressilator with multiple phase-separating TFs could generate oscillations with lower fluctuations in amplitude. Our framework, thus, enriches the synthetic biologists” toolkit for constructing robust and noise-tolerant circuits.

Several avenues remain for future investigation. Our model connected regulatory motifs with binary partitioning of TFs, whereas phase separation in cellular setting often involves multicomponent interactions, nonequilibrium aspects, and coupling to extrinsic noise [47, 55–57]. It would be intriguing to extend this framework in these directions to gain insights into various regulatory processes in cells. Additionally, connecting this framework to information-theoretic principles of regulation could reveal the governing informational roles that emerge due to phase separation of TFs associated with gene regulatory networks.

## ACKNOWLEDGEMENT

SC acknowledges the support provided by the DBT Ramalingaswami Fellowship.

## Supporting information

## S1 Phase separation in a simple binary mixture

We consider a minimal model of phase separation in a binary mixture, where transcription factor (TF) molecules– present at a fixed concentration–constitute one of the components (Fig. S1) [1, 2, 3]. We note that we adopt this binary mixture model following the formulation of Klosin et al. [1]. In this framework, the total system volume *V* is partitioned into two phases, a dense droplet phase of volume *V*_−_ and a surrounding dilute phase with volume *V*_+_ = *V* − *V*_−_. The total number of TF molecules is denoted by *p*, of which *p*_+_ resides in the dilute phase, while the remaining *p*_−_ = *p* − *p*_+_ are sequestered into the droplet phase. We now define effective volume fractions for the two phases as *ϕ*_*±*_ = *vp*_*±*_*/V*_*±*_, with *v* is the molecular volume of a single TF.

We assume that the droplet phase is composed entirely of TFs, with no solvent content. This leads to the volume fraction *ϕ*_−_ = 1. However, the dilute phase contains both TFs and solvent, leading to a volume fraction *ϕ*_+_ *<* 1 (Fig. S1). The total free energy of this system is given by,

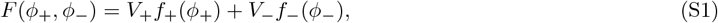

where f_+_(*ϕ*_+_) and *f*_−_(*ϕ*_−_) denote the free energy densities of the dilute and droplet phases, respectively. Under the assumption *ϕ*_−_ = 1, the free energy density of the droplet phase is approximated by *f*_−_(*ϕ*_−_) = − *µ/v*, where *µ* is the relative chemical potential. The free energy density of the dilute phase follows the ideal mixing form, *f*_+_(*ϕ*_+_) = (*k*_*B*_*T/v*) (*ϕ*_+_ ln *ϕ*_+_ − *ϕ*_+_), where *k*_*B*_ and *T* stand for Boltzmann constant and absolute temperature, respectively. Substituting these expressions into Eq. (S1), the total free energy becomes, *F* (*ϕ*_+_, *ϕ*_−_) = *k*_*B*_*T* (*V*_+_*/v*) (*ϕ*_+_ ln *ϕ*_+_ − *ϕ*_+_) − *µ*(*V*_−_*/v*). However, the total free energy can also be expressed in terms of number of TFs in the two phases as,

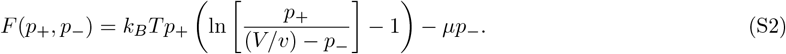

While writing Eq. (S2), we use *V*_−_*/v* = *p*_−_, which follows from *ϕ*_−_ = 1. Rewriting this equation as the functions of *p*_+_ and *p*, we have,

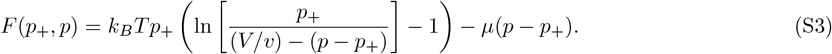

We now define the volume fraction of the whole system by *ϕ* = *vp/V*. After some algebraic simplification, Eq. (S3) can be rewritten in terms of *ϕ*_+_ and *ϕ* as,

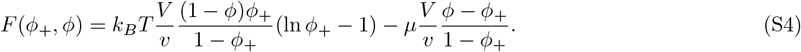

To determine the condition under which phase separation occurs, we minimize the total free energy with respect to the dilute-phase volume fraction *ϕ*_+_, while keeping the total TF volume fraction *ϕ* fixed. The solution to this variational condition 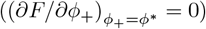 determines the threshold volume fraction, *ϕ*^∗^, at which phase separation is initiated. This yields,

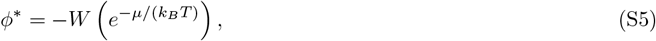

**Figure S1:**
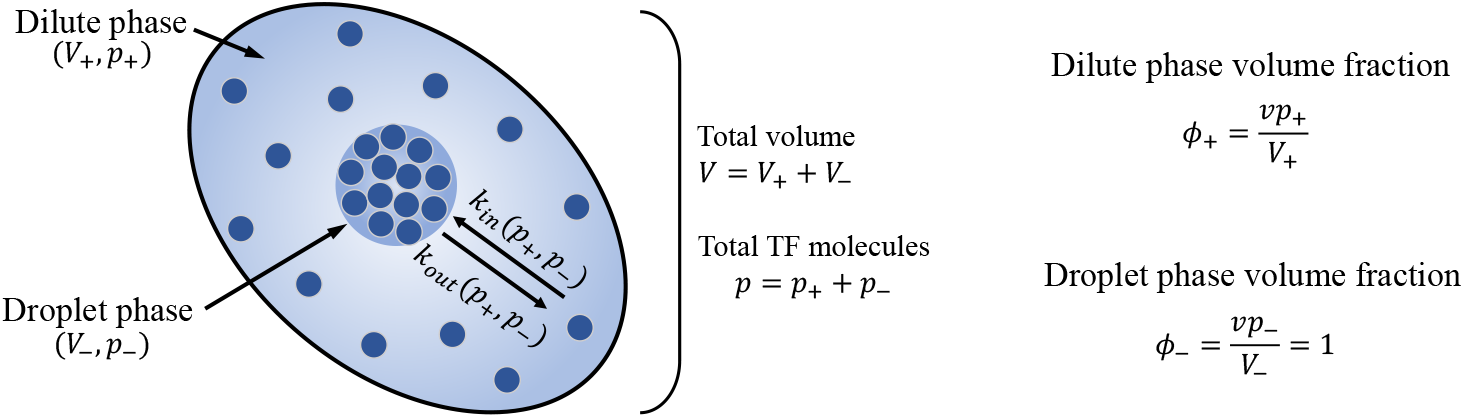
Phase separation of TFs in a binary mixture. The total TF population is partitioned between two coexisting phases: a dense droplet phase of volume *V*_−_ containing *p*_−_ TF molecules, and a surrounding dilute phase of volume *V*_+_ = *V* − *V*_−_ containing *p*_+_ TF molecules. It is assumed that the droplet phase is filled with TFs only and no solvent content is present, yielding the volume fraction *ϕ*_−_ = 1. On the contrary, the volume fraction in the dilute phase satisfies *ϕ*_+_ *<* 1.

where, *W* () denotes the Lambert W function. This threshold defines the critical volume fraction of TFs in the dilute phase above which it becomes thermodynamically favorable for the system to separate into coexisting dense and dilute phases. By using Eq. (S5), we can determine the value of *µ* corresponding to a given *ϕ*^∗^. Since *ϕ*^∗^ is related to the threshold TF copy number *p*^∗^ by *ϕ*^∗^ = *vp*^∗^*/V*, specifying a value for *p*^∗^ directly sets the chemical potential *µ* that governs the onset of phase separation. In the present study, we treat *p*^∗^ as the primary tuning parameter to control the initiation of phase separation. Using this definition of *ϕ*^∗^, we express the chemical potential as,

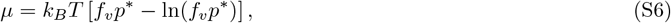

where *f*_*v*_ = *v/V* is the volume factor. This expression of *µ*, determined by the threshold TF copy number *p*^∗^, is then used in Eq. (S2) to compute the total free energy of the system for a given *p*_+_ and *p*_−_.

We now consider the stochastic exchange of TFs between the dilute and droplet phases, assuming a fixed total copy number *p* (Fig. S1). The transition rate from the dilute to droplet phase is denoted by *k*_*in*_(*p*_+_, *p*_−_), while the reverse transition rate is given by *k*_*out*_(*p*_+_, *p*_−_). At equilibrium, this exchange satisfies the microscopic detailed balance condition, expressed as,

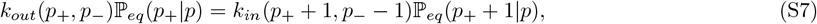

where ℙ_*eq*_(*p*_+_|*p*) denotes the equilibrium probability of observing *p*_+_ TFs in the dilute phase, with the droplet phase containing the remaining *p*_−_ = *p* − *p*_+_ molecules. This probability follows the Boltzmann distribution, ℙ_*eq*_(*p*_+_|*p*) ∝ exp [−*F* (*p*_+_, *p*_−_)*/*(*k*_*B*_*T*)]. Using this relation in Eq. (S7), we can express the transition rate *k*_*out*_ in term of *k*_*in*_ as

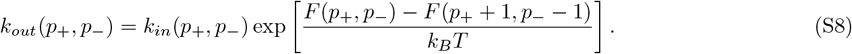

The transition rate *k*_*in*_ is assumed to be diffusion-limited [1], and is given by,

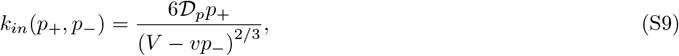

where, the diffusion coefficient 𝒟_*p*_ = *V* ^2*/*3^*/*(6*τ*_*D*_). Here *τ*_*D*_ denotes the characteristic diffusion time for a single TF molecule across the system volume *V*. We note that this diffusion time sets the time scale of phase separation. To compare diffusion with TF turnover, we introduce the dimensionless ratio (a scaling parameter) *ω*_*p*_ = *τ*_*D*_*/τ*_*p*_, where *τ*_*p*_ is the characteristic timescale for TF degradation. This parameter becomes relevant when we consider TF dynamics explicitly. Now, expressing the diffusion constant in terms of *ω*_*p*_, we write,

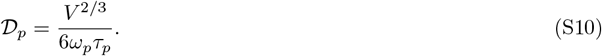

The scaling parameter *ω*_*p*_ quantifies the competition between diffusion and degradation. When *ω*_*p*_ *<* 1, diffusion dominates, enabling phase separation to influence TF dynamics. In contrast, *ω*_*p*_ *>* 1 implies rapid degradation of the TFs, limiting the impact of phase separation. This ratio thus determines the extent to which phase separation can modulate gene regulatory behavior, as explored in Sec S2. The parameters for phase separation, which are common to both toggle switch and repressilator are given in Table S1.

## S2 Role of timescale separation between diffusion and degradation in regulating circuit dynamics

In this section, we examine how the scaling parameter *ω*_*p*_ regulates the mean-field dynamics of the TFs in both the toggle switch and repressilator. The scaling parameter quantifies the timescale separation between the TF diffusion to the droplet phase and the TF degradation. A smaller value of *ω*_*p*_, which corresponds to shorter diffusion times relative to the degradation timescale, yields a larger effective diffusion coefficient 𝒟_*p*_. This accelerates droplet formation and thereby more strongly perturbs the dynamics of both motifs. Conversely, a larger *ω*_*p*_ slows droplet formation, reducing the overall influence of phase separation on the system”s dynamics. We quantitatively analyze the effect of *ω*_*p*_ in the following.

### Toggle switch

To explore the effect of timescale competition between diffusion and TF degradation, we vary the scaling parameter *ω*_*p*_ for phase-separating TFs under different phase-separation scenarios. As a representative case, we first consider Y to undergo phase separation, with the threshold copy number for phase separation set at *y*^∗^ = 50. The corresponding scaling parameter is defined as *ω*_*y*_ = *τ*_*d*_*/τ*_*y*_, where *τ*_*y*_ denotes the degradation timescale of Y (see Table S2). This definition allows us to compute the diffusion coefficient 𝒟_*y*_ from Eq. (S10). We find that increasing *ω*_*y*_ leads to an upward shift of the fixed point {*x=low, y=high*} along the X-nullcline (Fig. S2a). Moreover, the curvature of the separatrix bending toward the basin {*x=low, y=high*} becomes less pronounced at higher *ω*_*y*_. These results indicate that the influence of phase separation on TF dynamics becomes more prominent when the diffusion time is shorter relative to the degradation timescale.

When both X and Y undergo phase separation, the relative values of their scaling parameters (*ω*_*x*_ and *ω*_*y*_) determine the dynamical landscape of the toggle switch (Fig. S2b). Using Eq. (S10), we compute the corresponding diffusion coefficients for both TFs, with the degradation timescales *τ*_*x*_ and *τ*_*y*_ taken from Table S2. When *ω*_*x*_ and *ω*_*y*_ differ, the TF with the shorter diffusion time dominates the dynamics, causing the separatrix to bend toward its associated basin of attraction. This results in an asymmetric basin structure that favours one stable state over the other. Conversely, when *ω*_*x*_ and *ω*_*y*_ are equal, the basin symmetry is restored.

### Repressilator

In the repressilator, we consider all the TFs–X, Y, and Z–to undergo phase separation with identical scaling parameters (*ω*_*x*_ = *ω*_*y*_ = *ω*_*z*_) for varying thresholds–*x*^∗^, *y*^∗^, and *z*^∗^. These parameters are varied simultaneously while maintaining equality, as shown in Fig. S2c. Changing this common timescale ratio produces distinct effects on the oscillation amplitude. When diffusion is fast compared to the degradation, phase separation clamps the oscillatory peaks near the threshold copy number, reflecting effective concentration buffering by the droplet phase. As diffusion becomes slower relative to TF degradation, this buffering effect weakens, and the oscillatory dynamics become increasingly governed by the intrinsic synthesis-degradation kinetics of the circuit. Consequently, the oscillation amplitude rises progressively as the system transitions from a diffusion-dominated to a reaction-dominated regime (Fig. S2c).

These findings reveal that the regulatory impact of phase separation is most pronounced for TFs that diffuse rapidly to the droplet phase and are degraded slowly. Fast diffusion enables such TFs to efficiently exchange between the dilute and droplet phases, allowing condensates to exert a stronger influence on their effective dynamics. At the same time, slower degradation prolongs their residence within the regulatory network, amplifying the cumulative effects of phase separation on circuit behavior.

**Figure S2:**
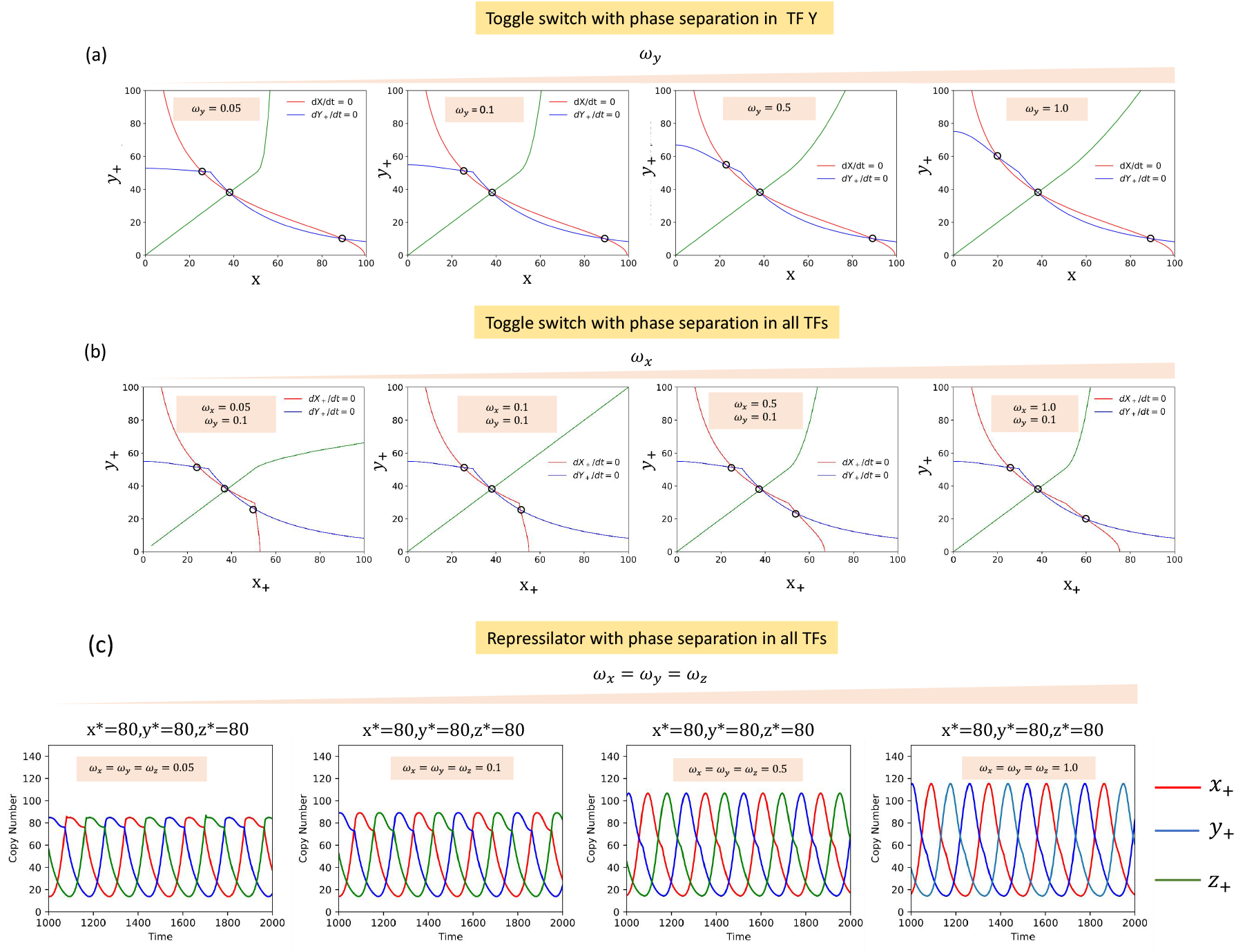
Effects of timescale separation between TF diffusion and degradation. (a) Toggle switch with phase separation in TF Y. (b) Toggle switch with phase separation in both TFs. (c) Repressilator with phase separation in all TFs. Green curves in panels a–b indicate the separatrix dividing the two basins of attraction; open circles on these curves mark unstable fixed points, while the remaining open circles correspond to stable fixed points. The parameters used to generate these plots are given in Tables S1-S3.

## S3 Stochastic modeling and noise quantification

We simulate the dynamics of the toggle switch and the repressilator under different phase-separation scenarios using the stochastic simulation algorithm (SSA). Each reaction is characterized by a corresponding propensity function (*a*_*i*_). The waiting time between successive reactions is sampled from an exponential distribution with mean 1*/a*_0_, where *a*_0_ = ∑_*i*_ *a*_*i*_. The next reaction is then selected by drawing a random number from a uniform distribution weighted by the cumulative propensities. After each selection, the molecular species are updated according to the stoichiometry of the chosen reaction, and the process is iterated until the desired final time is reached.

**Figure S3:**
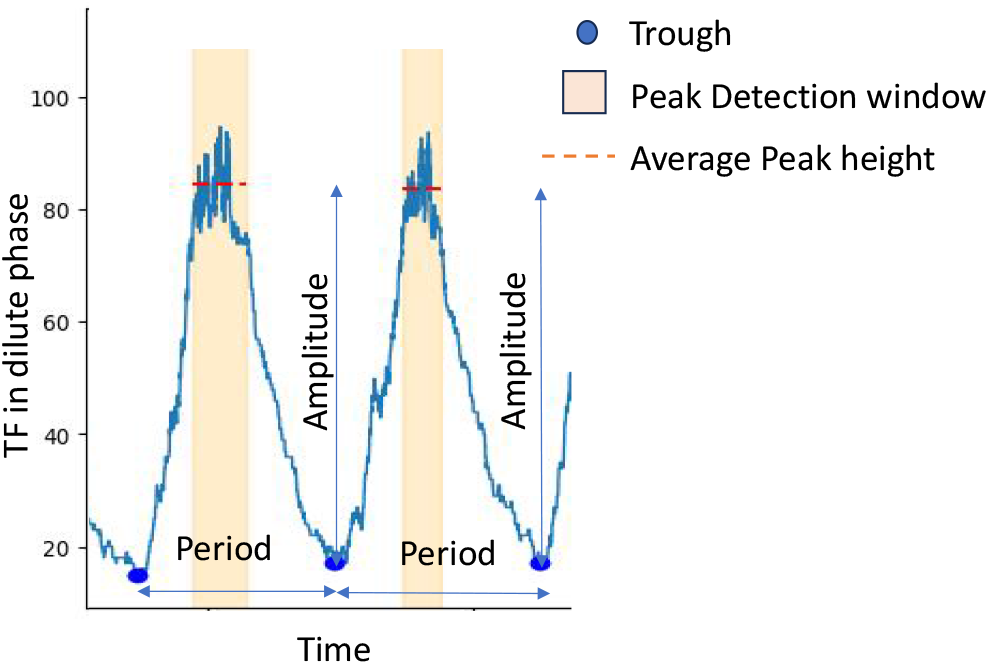
A schematic of detecting oscillation amplitude and period from the simulated trajectory of TF in the repressilator.

The reaction schemes and their corresponding propensity functions for the canonical toggle switch and repressilator are listed in Tables S4 and S7, respectively. To incorporate phase separation of TFs, each TF is considered to be synthesized in the dilute phase. When its copy number exceeds a threshold 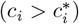, transitions between the dilute and droplet phases occur with thermodynamically defined rates *k*_*in*_ and *k*_*out*_. The reaction schemes and the propensity functions of the toggle switch with phase separation in TF Y, and in both TFs are provided in Tables S5 and S6, respectively. Similarly, the corresponding reaction schemes for the repressilator with phase separation in TF Z, in TFs Y and Z, and in all three TFs are given in Tables S8, S9, and S10, respectively.

The stochastic trajectories generated by the SSA are used to construct the distributions of TF abundances in the two stable states of the toggle switch, and the distributions of oscillation amplitude and period in the repressilator, for both canonical and phase-separating cases. The method for quantifying oscillatory features in the repressilator is described below. The oscillation period is defined as the time interval between two successive troughs in the stochastic trajectory (see Fig. S3). Between each pair of troughs, a local maximum is identified, and the peak value is computed by averaging the trajectory values within a window centered around 80% of that maximum. The oscillation amplitude is then determined as the difference between this averaged peak value and the subsequent trough.

For the toggle switch, the steady-state distributions of TF copy numbers provide the mean (⟨*c*_*i*_⟩) and standard deviation 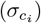 for each TF. The noise is quantified using the coefficient of variation (CV), defined as 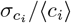. For the repressilator, the extracted distributions of oscillation amplitude and period are analyzed similarly. The noise in these quantities is quantified using their respective CVs, CV_Amplitude_ = *σ*_Amplitude_*/*⟨Amplitude⟩ and CV_Period_ = *σ*_Period_*/* ⟨Period⟩. These measures capture the relative magnitude of fluctuations around the steady state and enable direct comparison across different phase separation scenarios.

**Table S1:** Model parameters for the binary phase separation model [1].

| Parameter | Notation | Value | Remarks |
| --- | --- | --- | --- |
| Boltzmann constant | $k_B$ | $1.38 \times 10^{-23} \text{ J K}^{-1}$ | Physical constant |
| Absolute temperature | $T$ | 305 K | Fixed physiological temperature |
| Total system volume | $V$ | $10^{-20} \text{ L}$ | Effective cellular volume |
| Molecular volume of a TF | $v$ | $10^{-25} \text{ L}$ | Assumed uniform for all TFs in toggle switch and repressilator |
| Diffusion–degradation time scale separation | $\omega_p$ | 0.1 | Common to all TFs in toggle switch and repressilator |

**Table S2:**
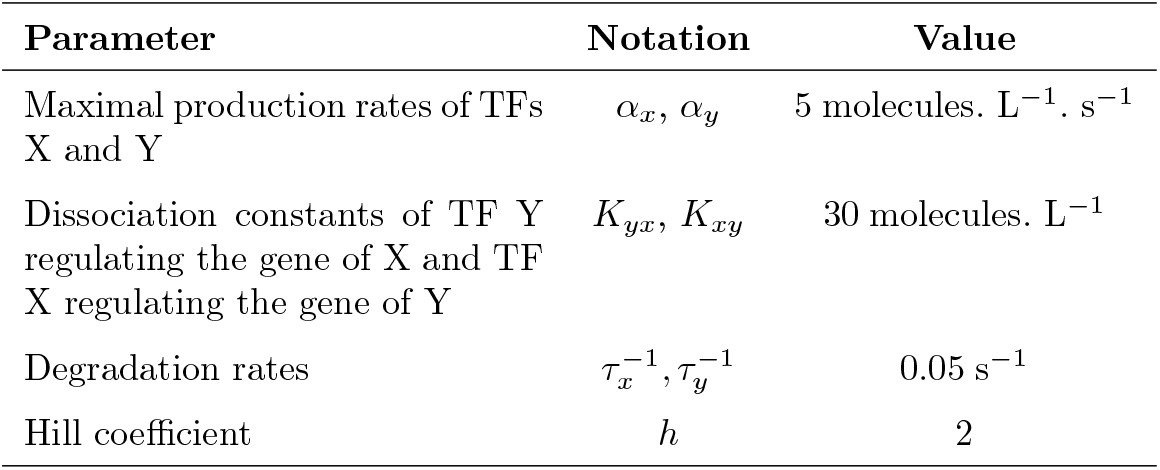
Model parameters for the canonical toggle switch.

**Table S3:** Model parameters for the canonical repressilator.

| Parameter | Notation | Value |
| --- | --- | --- |
| Maximal production rates of TF X, TF Y, TF Z | $\alpha_x, \alpha_y, \alpha_z$ | 5 molecules. $\text{L}^{-1} \cdot \text{s}^{-1}$ |
| Dissociation constants of TF Z regulating the gene of X, TF X regulating the gene of Y, TF Y regulating the gene of Z | $K_{zx}, K_{xy}, K_{yz}$ | 20 molecules. $\text{L}^{-1}$ |
| Degradation rates of X, Y, Z | $\tau_x^{-1}, \tau_y^{-1}, \tau_z^{-1}$ | $0.02 \text{ s}^{-1}$ |
| Hill coefficient | $h$ | 3 |

**Table S4:** Reaction schemes and corresponding propensities for the canonical toggle switch.

| Description | Reaction step | Propensity |
| --- | --- | --- |
| Synthesis of TF X | $x \rightarrow x + 1$ | $\frac{\alpha_x}{1 + (y/K_{yx})^h}$ |
| Degradation of TF X | $x \rightarrow x - 1$ | $\tau_x^{-1} x$ |
| Synthesis of TF Y | $y \rightarrow y + 1$ | $\frac{\alpha_y}{1 + (x/K_{xy})^h}$ |
| Degradation of TF Y | $y \rightarrow y - 1$ | $\tau_y^{-1} y$ |

**Table S5:** Reaction schemes and corresponding propensities for the toggle switch with phase separation in TF Y.

| Description | Reaction step | Propensity |
| --- | --- | --- |
| Synthesis of TF X | $x \rightarrow x + 1$ | $\frac{\alpha_x}{1+(y_+/K_{yx})^h}$ |
| Degradation of TF X | $x \rightarrow x - 1$ | $\tau_x^{-1} x$ |
| Synthesis of TF Y in dilute phase | $y_+ \rightarrow y_+ + 1$ | $\frac{\alpha_y}{1+(x/K_{xy})^h}$ |
| Degradation of TF Y in dilute phase | $y_+ \rightarrow y_+ - 1$ | $\tau_y^{-1} y_+$ |
| Transfer of TF Y from dilute to droplet phase (if $y_+ > y^*$) | $y_+ \rightarrow y_+ - 1, y_- \rightarrow y_- + 1$ | $k_{in}(y_+, y_-)$ |
| Transfer of TF Y from droplet to dilute phase | $y_+ \rightarrow y_+ + 1, y_- \rightarrow y_- - 1$ | $k_{out}(y_+, y_-)$ |
| Degradation of TF Y in droplet phase | $y_- \rightarrow y_- - 1$ | $\tau_y^{-1} y_-$ |

**Table S6:** Reaction schemes and corresponding propensities for the toggle switch with phase separation in both TFs.

| Description | Reaction step | Propensity |
| --- | --- | --- |
| Synthesis of TF X in dilute phase | $x_+ \rightarrow x_+ + 1$ | $\frac{\alpha_x}{1+(y_+/K_{yx})^h}$ |
| Degradation of TF X in dilute phase | $x_+ \rightarrow x_+ - 1$ | $\tau_x^{-1} x_+$ |
| Transfer of TF X from dilute to droplet phase (if $x_+ > x^*$) | $x_+ \rightarrow x_+ - 1, x_- \rightarrow x_- + 1$ | $k_{in}(x_+, x_-)$ |
| Transfer of TF X from droplet to dilute phase | $x_+ \rightarrow x_+ + 1, x_- \rightarrow x_- - 1$ | $k_{out}(x_+, x_-)$ |
| Degradation of TF X in droplet phase | $x_- \rightarrow x_- - 1$ | $\tau_x^{-1} x_-$ |
| Synthesis of TF Y in dilute phase | $y_+ \rightarrow y_+ + 1$ | $\frac{\alpha_y}{1+(x_+/K_{xy})^h}$ |
| Degradation of TF Y in dilute phase | $y_+ \rightarrow y_+ - 1$ | $\tau_y^{-1} y_+$ |
| Transfer of TF Y from dilute to droplet phase (if $y_+ > y^*$) | $y_+ \rightarrow y_+ - 1, y_- \rightarrow y_- + 1$ | $k_{in}(y_+, y_-)$ |
| Transfer of TF Y from droplet to dilute phase | $y_+ \rightarrow y_+ + 1, y_- \rightarrow y_- - 1$ | $k_{out}(y_+, y_-)$ |
| Degradation of TF Y in droplet phase | $y_- \rightarrow y_- - 1$ | $\tau_y^{-1} y_-$ |

**Table S7:**
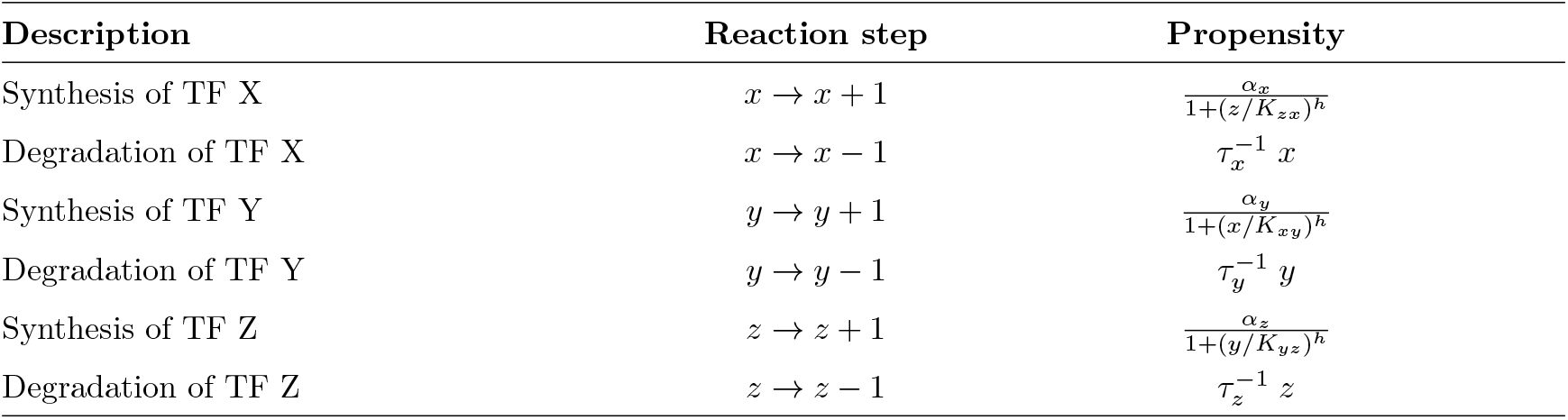
Reaction schemes and corresponding propensities for the canonical repressilator.

**Table S8:**
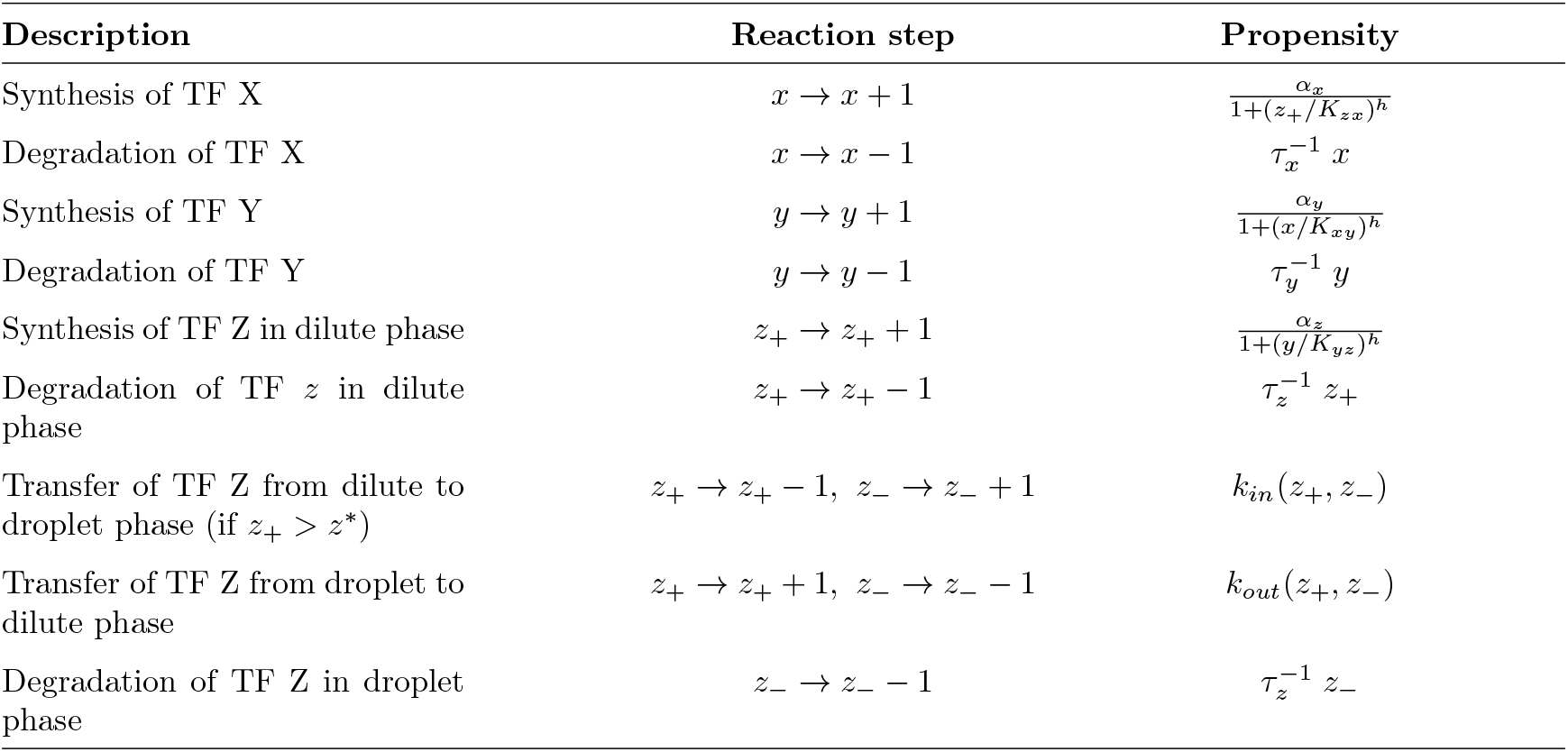
Reaction schemes and corresponding propensities for the repressilator with phase separation in TF Z.

| Description | Reaction step | Propensity |
| --- | --- | --- |
| Synthesis of TF X | $x \rightarrow x + 1$ | $\frac{\alpha_x}{1+(z_+/K_{zx})^h}$ |
| Degradation of TF X | $x \rightarrow x - 1$ | $\tau_x^{-1} x$ |
| Synthesis of TF Y | $y \rightarrow y + 1$ | $\frac{\alpha_y}{1+(x/K_{xy})^h}$ |
| Degradation of TF Y | $y \rightarrow y - 1$ | $\tau_y^{-1} y$ |
| Synthesis of TF Z in dilute phase | $z_+ \rightarrow z_+ + 1$ | $\frac{\alpha_z}{1+(y/K_{yz})^h}$ |
| Degradation of TF $z$ in dilute phase | $z_+ \rightarrow z_+ - 1$ | $\tau_z^{-1} z_+$ |
| Transfer of TF Z from dilute to droplet phase (if $z_+ > z^*$) | $z_+ \rightarrow z_+ - 1, z_- \rightarrow z_- + 1$ | $k_{in}(z_+, z_-)$ |
| Transfer of TF Z from droplet to dilute phase | $z_+ \rightarrow z_+ + 1, z_- \rightarrow z_- - 1$ | $k_{out}(z_+, z_-)$ |
| Degradation of TF Z in droplet phase | $z_- \rightarrow z_- - 1$ | $\tau_z^{-1} z_-$ |

**Table S9:** Reaction schemes and corresponding propensities for the repressilator with phase separation in TFs Y and Z.

| Description | Reaction step | Propensity |
| --- | --- | --- |
| Synthesis of TF X | $x \rightarrow x + 1$ | $\frac{\alpha_x}{1+(z_+/K_{zx})^h}$ |
| Degradation of TF X | $x \rightarrow x - 1$ | $\tau_x^{-1} x$ |
| Synthesis of TF Y in dilute phase | $y_+ \rightarrow y_+ + 1$ | $\frac{\alpha_y}{1+(x/K_{xy})^h}$ |
| Degradation of TF Y in dilute phase | $y_+ \rightarrow y_+ - 1$ | $\tau_y^{-1} y_+$ |
| Transfer of TF Y from dilute to droplet phase (if $y_+ > y^*$) | $y_+ \rightarrow y_+ - 1, y_- \rightarrow y_- + 1$ | $k_{in}(y_+, y_-)$ |
| Transfer of TF Y from droplet to dilute phase | $y_+ \rightarrow y_+ + 1, y_- \rightarrow y_- - 1$ | $k_{out}(y_+, y_-)$ |
| Degradation of TF Y in droplet phase | $y_- \rightarrow y_- - 1$ | $\tau_y^{-1} y_-$ |
| Synthesis of TF Z in dilute phase | $z_+ \rightarrow z_+ + 1$ | $\frac{\alpha_z}{1+(y_+/K_{yz})^h}$ |
| Degradation of TF Z in dilute phase | $z_+ \rightarrow z_+ - 1$ | $\tau_z^{-1} z_+$ |
| Transfer of TF Z from dilute to droplet phase (if $z_+ > z^*$) | $z_+ \rightarrow z_+ - 1, z_- \rightarrow z_- + 1$ | $k_{in}(z_+, z_-)$ |
| Transfer of TF Z from droplet to dilute phase | $z_+ \rightarrow z_+ + 1, z_- \rightarrow z_- - 1$ | $k_{out}(z_+, z_-)$ |
| Degradation of TF Z in droplet phase | $z_- \rightarrow z_- - 1$ | $\tau_z^{-1} z_-$ |

**Table S10:** Reaction schemes and corresponding propensities for the repressilator with phase separation in all TFs.

| Description | Reaction step | Propensity |
| --- | --- | --- |
| Synthesis of TF X in dilute phase | $x_+ \rightarrow x_+ + 1$ | $\frac{\alpha_x}{1+(z_+/K_{zx})^h}$ |
| Degradation of TF X in dilute phase | $x_+ \rightarrow x_+ - 1$ | $\tau_x^{-1} x_+$ |
| Transfer of TF X from dilute to droplet phase (if $x_+ > x^*$) | $x_+ \rightarrow x_+ - 1, x_- \rightarrow x_- + 1$ | $k_{in}(x_+, x_-)$ |
| Transfer of TF X from droplet to dilute phase | $x_+ \rightarrow x_+ + 1, x_- \rightarrow x_- - 1$ | $k_{out}(x_+, x_-)$ |
| Degradation of TF X in droplet phase | $x_- \rightarrow x_- - 1$ | $\tau_x^{-1} x_-$ |
| Synthesis of TF Y in dilute phase | $y_+ \rightarrow y_+ + 1$ | $\frac{\alpha_y}{1+(x_+/K_{xy})^h}$ |
| Degradation of TF Y in dilute phase | $y_+ \rightarrow y_+ - 1$ | $\tau_y^{-1} y_+$ |
| Transfer of TF Y from dilute to droplet phase (if $y_+ > y^*$) | $y_+ \rightarrow y_+ - 1, y_- \rightarrow y_- + 1$ | $k_{in}(y_+, y_-)$ |
| Transfer of TF Y from droplet to dilute phase | $y_+ \rightarrow y_+ + 1, y_- \rightarrow y_- - 1$ | $k_{out}(y_+, y_-)$ |
| Degradation of TF Y in droplet phase | $y_- \rightarrow y_- - 1$ | $\tau_y^{-1} y_-$ |
| Synthesis of TF Z in dilute phase | $z_+ \rightarrow z_+ + 1$ | $\frac{\alpha_z}{1+(y_+/K_{yz})^h}$ |
| Degradation of TF Z in dilute phase | $z_+ \rightarrow z_+ - 1$ | $\tau_z^{-1} z_+$ |
| Transfer of TF Z from dilute to droplet phase (if $z_+ > z^*$) | $z_+ \rightarrow z_+ - 1, z_- \rightarrow z_- + 1$ | $k_{in}(z_+, z_-)$ |
| Transfer of TF z from droplet to dilute phase | $z_+ \rightarrow z_+ + 1, z_- \rightarrow z_- - 1$ | $k_{out}(z_+, z_-)$ |
| Degradation of TF Z in droplet phase | $z_- \rightarrow z_- - 1$ | $\tau_z^{-1} z_-$ |

## References

[1] E. H. Davidson, The Regulatory Genome: Gene Regulatory Networks in Development and Evolution (Academic Press, Burlington, MA, 2006).

[2] U. Alon, An Introduction to Systems Biology: Design Principles of Biological Circuits (CRC Press, Boca Raton, FL, 2006).

[3] M. Kashima, Y. Shida, T. Yamashiro, H. Hirata, and H. Kurosaka, Intracellular and intercellular gene regulatory network inference from time-course individual rna-seq, Front. Bioinform. 1, 777299 (2021).

[4] S. S. Shen-Orr, R. Milo, S. Mangan, and U. Alon, Network motifs in the transcriptional regulation network of Escherichia coli, Nat. Genet. 31, 64 (2002).

[5] S. Mangan and U. Alon, Structure and function of the feed-forward loop network motif, Proc. Natl. Acad. Sci. U.S.A. 100, 11980 (2003).

[6] W. A. Lim, C. M. Lee, and C. Tang, Design principles of regulatory networks: Searching for the molecular algorithms of the cell, Mol. cell 49, 202 (2013).

[7] T. S. Gardner, C. R. Cantor, and J. J. Collins, Construction of a genetic toggle switch in Escherichia coli, Nature 403, 339 (2000).

[8] M. B. Elowitz and S. Leibler, A synthetic oscillatory network of transcriptional regulators, Nature 403, 335 (2000).

[9] S. M. Lyons, W. Xu, J. Medford, and A. Prasad, Loads bias genetic and signaling switches in synthetic and natural systems, Plos Comput. Biol. 10, e1003533 (2014).

[10] P. E. M. Purnick and R. Weiss, The second wave of synthetic biology: from modules to systems, Nat. Rev. Mol. Cell Biol. 10, 410 (2009).

[11] A. A. K. Nielsen and C. A. Voigt, Multi-input crispr/cas genetic circuits that interface host regulatory networks, Mol. Syst. Biol. 10, 763 (2014).

[12] J. Pomerening, E. Sontag, and J. Ferrell, Building a cell cycle oscillator: hysteresis and bistability in the activation of cdc2, Nat. Cell Biol. 5, 346 (2003).

[13] A. Arkin, J. Ross, and H. H. McAdams, Stochastic kinetic analysis of developmental pathway bifurcation in phage -infected escherichia coli cells, Genetics 149, 1633 (1998).

[14] T. Tian and K. Burrage, Stochastic models for regulatory networks of the genetic toggle switch, Proc. Natl. Acad. Sci. 103, 8372 (2006).

[15] E. M. Ozbudak, M. Thattai, H. N. Lim, B. I. Shraiman, and A. van Oudenaarden, Multistability in the lactose utilization network of escherichia coli, Nature 427, 737–740 (2004).

[16] M. Santillán, M. C. Mackey, and E. S. Zeron, Origin of bistability in the lac operon, Biophys. J. 92, 3830 (2007).

[17] C. P. Bagowski and J. Ferrell, Bistability in the jnk cascade, Curr. Biol. 11, 1176 (2001).

[18] W. Xiong and J. Ferrell, A positive-feedback-based bistable “memory module” that governs a cell fate decision, Nature 426, 460 (2003).

[19] N. I. Markevich, J. B. Hoek, and B. N. Kholodenko, Signaling switches and bistability arising from multisite phosphorylation in protein kinase cascades, J. Cell Biol. 164, 353 (2004).

[20] A. Hardig, T. Tian, E. Westbury, E. Frische, and J. F. Hancock, Subcellular localization determines map kinase signal output, CUrr. Biol. 15, 869 (2005).

[21] M. R. Doyle, S. J. Davis, R. M. Bastow, H. G. McWatters, L. Kozma-Bognár, F. Nagy, A. J. Millar, and R. M. Amasino, The elf4 gene controls circadian rhythms and flowering time in arabidopsis thaliana, Nature 419, 74 (2002).

[22] J. Garcia-Ojalvo, M. B. Elowitz, and S. H. Strogatz, Modeling a synthetic multicellular clock: Repressilators coupled by quorum sensing, Proc. Natl. Acad. Sci. 101, 10955 (2004).

[23] D. H. Nagel and S. A. Kay, Complexity in the wiring and regulation of plant circadian networks, Curr. Biol. 22, R648 (2012).

[24] A. Pokhilko, A. P. Fernández, K. D. Edwards, M. M. Southern, K. J. Halliday, and A. J. Millar, The clock gene circuit in <i>arabidopsis</i> includes a repressilator with additional feedback loops, Mol. Syst. Biol. 8, 574 (2012).

[25] L. Wu, Q. Ouyang, and H. Wang, Robust network topologies for generating oscillations with temperature-independent periods, PLOS ONE 12, 1 (2017).

[26] Q. l. DOng, X. y. Xing, Y. Han, X. l. Wei, and S. Zhang, De novo organelle biogenesis in the cyanobacterium tdx16 released from the green alga haematococcus pluvialis, CellBio. 9, 29 (2020).

[27] T. L. Tootle and I. Rebay, Post-translational modifications influence transcription factor activity: a view from the ets superfamily, Bioessays 27, 285 (2005).

[28] G. Balázsi, A. van Oudenaarden, and J. J. Collins, Cellular decision making and biological noise: From microbes to mammals, Cell 144, 910 (2011).

[29] C. Vogel and E. M. Marcotte, Insights into the regulation of protein abundance from proteomic and transcriptomic analyses, Nat. Rev. Genet. 13, 227 (2012).

[30] R. K. Yadav, A. S. Chauhan, L. Zhuang, and B. Gan, Foxo transcription factors in cancer metabolism, Sem. Cancer Biol. 50, 65 (2018).

[31] Q. Guo, Y. Jin, X. Chen, X. Ye, X. Shen, M. Lin, C. Zeng, T. Zhou, and J. Zhang, Nf-κb in biology and targeted therapy: new insights and translational implications, Sig. Transduct. Target Ther. 9, 53 (2024).

[32] M. Scott, C. W. Gunderson, E. M. Mateescu, Z. Zhang, and T. Hwa, Interdependence of cell growth and gene expression: Origins and consequences, Science 330, 1099 (2010).

[33] A. Y. Weiße, D. A. Oyarzún, V. Danos, and P. S. Swain, Mechanistic links between cellular trade-offs, gene expression, and growth, Proc. Natl. Acad. Sci. 112, E1038 (2015).

[34] J. M. Schmiedel, S. L. Klemm, Y. Zheng, A. Sahay, N. Blüthgen, D. S. Marks, and A. van Oudenaarden, Microrna control of protein expression noise, Science 348, 128 (2015).

[35] R. Sabi and T. Tuller, Modelling and measuring intracellular competition for finite resources during gene expression, J. R. Soc. Interface 16, 20180887 (2019).

[36] A. Burger, A. M. Walczak, and P. G. Wolynes, Influence of decoys on the noise and dynamics of gene expression, Phys. Rev. E 86, 041920 (2012).

[37] P. Bokes and A. Singh, Protein copy number distributions for a self-regulating gene in the presence of decoy binding sites, PLOS ONE 10, 1 (2015).

[38] M. Soltani, P. Bokes, Z. Fox, and A. Singh, Nonspecific transcription factor binding can reduce noise in the expression of downstream proteins, Phys. Biol. 12, 055002 (2015).

[39] S. Das and S. Choubey, Tunability enhancement of gene regulatory motifs through competition for regulatory protein resources, Phys. Rev. E 102, 052410 (2020).

[40] A. A. Hyman, C. A. Weber, and F. Jülicher, Liquid-liquid phase separation in biology, Annu. Rev. Cell Dev. Biol. 30, 39 (2014).

[41] S. F. Banani, H. O. Lee, A. A. Hyman, and M. K. Rosen, Biomolecular condensates: organizers of cellular biochemistry, Nat. Rev. Mol. Cell Biol. 18, 285 (2017).

[42] B. R. Sabari, A. Dall”Agnese, A. Boija, I. A. Klein, E. L. Coffey, K. Shrinivas, B. J. Abraham, N. M. Hannett, A. V. Zamudio, J. C. Manteiga, C. H. Li, Y. E. Guo, D. S. Day, J. Schuijers, E. Vasile, S. Malik, D. Hnisz, T. I. Lee, I. I. Cisse, R. G. Roeder, P. A. Sharp,A. K. Chakraborty, and R. A. Young, Coactivator condensation at super-enhancers links phase separation and gene control, Science 361, eaar3958 (2018).

[43] Boija, I. A. Klein, B. R. Sabari, A. Dall”Agnese, E. L. Coffey, A. V. Zamudio, C. H. Li, K. Shrinivas, J. C. Manteiga, N. M. Hannett, J. Abraham, L. K. Afeyan, Y. E. Guo, J. K. Rimel, C. B. Fant, J. Schuijers, T. I. Lee, D. J. Taatjes, and R. A. Young, Transcription factors activate genes through the phase-separation capacity of their activation domains, Cell 175, 1842 (2018).

[44] J. Plys and R. E. Kingston, Dynamic condensates activate transcription, Science 361, 329 (2018).

[45] Hnisz, K. Shrinivas, R. A. Young, A. K. Chakraborty, and P. A. Sharp, A phase separation model for transcriptional control, Cell 169, 13 (2017).

[46] W.-K. Cho, J.-H. Spille, M. Hecht, C. Lee, C. Li, V. Grube, and I. I. Cisse, Mediator and rna polymerase ii clusters associate in transcription-dependent condensates, Science 361, 412 (2018).

[47] A. Klosin, F. Oltsch, T. Harmon, A. Honigmann, F. Jülicher, A. A. Hyman, and C. Zechner, Phase separation provides a mechanism to reduce noise in cells, Science 367, 464 (2020).

[48] R. Zhang, W. Yang, R. Zhang, S. Rijal, A. Youssef, W. Zheng, and X.-J. Tian, Phase separation to resolve growth-related circuit failures, bioRxiv 10.1101/2024.11.01.621586 (2024).

[49] L. Hong, Z. Wang, Z. Zhang, S. Luo, T. Zhou, and J. Zhang, Phase separation reduces cell-to-cell variability of transcriptional bursting, Math. Biosci. 367, 109127 (2024).

[50] L. Hong, Z. Zhang, Z. Wang, X. Yu, and J. Zhang, Phase separation provides a mechanism to drive phenotype switching, Phys. Rev. E 109, 064414 (2024).

[51] T. Gillespie, A general method for numerically simulating the stochastic time evolution of coupled chemical reactions, J. Comp. Phys. 22, 403 (1976).

[52] T. Gillespie, Exact stochastic simulation of coupled chemical reactions, J. Phys. Chem. 81, 2340 (1977).

[53] D.-D. C. L. Z. D. P. D. G. M. C. C. H. A. B. S. L. L. D. X. Chong, S. and R. Tjian, Imaging dynamic and selective low-complexity domain interactions that control gene transcription, Science 361, eaar2555 (2018).

[54] M. B. Elowitz, A. J. Levine, E. D. Siggia, and P. S. Swain, Stochastic gene expression in a single cell, Science 297, 1183 (2002).

[55] A. Patel, H. O. Lee, L. Jawerth, S. Maharana, M. Jahnel, M. Y. Hein, S. Stoynov, J. Mahamid, S. Saha, T. M. Franzmann, A. Pozniakovski, Poser, N. Maghelli, L. A. Royer, M. Weigert, E. W. Myers, S. Grill, D. Drechsel, A. A. Hyman, and S. Alberti, A liquid-to-solid phase transition of the als protein fus accelerated by disease mutation, Cell 162, 1066 (2015).

[56] S. Alberti, A. Gladfelter, and T. Mittag, Considerations and challenges in studying liquid-liquid phase separation and biomolecular condensates, Cell 176, 419 (2019).

[57] Zechner and F. Jülicher, Concentration buffering and noise reduction in non-equilibrium phase-separating systems, Cell Systems 16, 101168 (2025).

## References

[1] Klosin, F. Oltsch, T. Harmon, A. Honigmann, F. Jülicher, A. A. Hyman, and C. Zechner. Phase separation provides a mechanism to reduce noise in cells. Science, 367(6476):464–468, 2020.

[2] L. Hong, Z. Wang, Z. Zhang, S. Luo, T. Zhou, and J. Zhang. Phase separation reduces cell-to-cell variability of transcriptional bursting. Math. Biosci., 367:109127, 2024.

[3] L. Hong, Z. Zhang, Z. Wang, X. Yu, and J. Zhang. Phase separation provides a mechanism to drive phenotype switching. Phys. Rev. E, 109:064414, 2024.

